# Niche evolution of the Neotropical tree genus *Otoba* in the context of global biogeography of the nutmeg family, Myristicaceae

**DOI:** 10.1101/2020.10.02.324368

**Authors:** Laura Frost, Daniel A. Santamaría-Aguilar, Daisy Singletary, Laura P. Lagomarsino

## Abstract

**Aim:** Plant distributions are influenced by species’ ability to colonize new areas via long-distance dispersal and propensity to adapt to new environments via niche evolution. We use *Otoba* (Myristicaceae), an ecologically dominant tree genus found in low-to-mid elevation wet forests, as a system to understand the relative importance of these processes within the Neotropics, a region characterized by high species richness and a diversity of biomes.

**Location:** Neotropics and global

**Taxon:** *Otoba* and entire Myristicaceae

**Methods:** We resolve the first phylogeny of *Otoba* using targeted sequence capture phylogenomics. We pair this with the most densely sampled phylogeny of Myristicaceae to date, inferred using publicly available data. We then use phylogenetic comparative methods to infer biogeography and examine patterns of niche evolution.

**Results:** Myristicaceae has an Old World origin, with a single expansion event into the Americas. Divergence dates, fossil evidence, and a notable lack of long-distance dispersal are consistent with a Boreotropical origin of Neotropical Myristicaceae. Mirroring the rarity of dispersal at the family level, *Otoba*’s biogeography is marked by few biogeographic events: two expansions into Central America from a South American ancestor and a single dispersal event across the Andes. This limited movement contrasts with rapid climatic niche evolution, typically occurring across geographically proximate habitats.

**Main conclusions:** Contrasting with previous studies, long-distance dispersal does not need to be invoked to explain the pantropical distribution of Myristicaceae, nor the biogeography of *Otoba*. This likely results from the family’s relatively large seeds that are dispersed by large-bodied vertebrates. Instead, rapid niche evolution in *Otoba* has facilitated its occurrence throughout mesic habitats of the northern Neotropics, including the Amazon rainforest and Andean montane forests. *Otoba* adds to a growing group of Neotropical plant clades in which climate adaptation following local migration is common, implying an important role of niche evolution in the assembly of the Neotropical flora.

**Significance statement:** Species distributions across the climatically and topographically heterogenous Neotropics are explained by a combination of local adaptation and dispersal. The relative importance of these mechanisms is clade dependent. We find that niche evolution in geographically proximal habitats is much more common than long-distance dispersal to preadapted regions in the tree genus *Otoba*, which includes both hyperdominant Amazonian species and narrow Andean endemics. The lack of long-distance dispersal is likely due to *Otoba*’s large seeds. Our results add to a growing body of literature demonstrating a key role of labile niche evolution across steep environmental gradients in Neotropical plant biogeography.

---

The distribution of plant clades is controlled by a combination of biogeographic movement, the pace of environmental adaptation, and macroevolutionary dynamics (Donoghue 2008). Movement into new regions is determined by geographic proximity and intrinsic traits that govern dispersal ability (e.g., propagule type), while adaptation to new habitats is a product of the degree of phylogenetic niche conservatism within a lineage (Edwards and Donoghue 2013). Mounting evidence suggests that the degree to which plant groups disperse to new locations to which they are pre-adapted versus shift their ecological tolerances to habitats in close geographic proximity is both clade and environment dependent. In some groups, like high Andean plant radiations derived from North American temperate clades (Pouchon et al. 2018; Hughes and Eastwood 2006; Madriñán, Cortés, and Richardson 2013) and mangroves (Woodroffe and Grindrod 1991), it is, to borrow a phrase from Donoghue (2008), “easier to move than evolve”. In other clades, including *Mimosa* and *Andira* (Fabaceae) in the Brazilian Cerrado (Simon et al. 2009), *Protium* (Burseraceae) on white sand soils in the Amazon and Guiana Shield (Fine and Baraloto 2016), and Montiaceae in temperate latitudes (Ogburn and Edwards 2015), major transitions in environmental tolerance are very common. In fact, a potentially large proportion of Neotropical biota is a result of migration between major biomes (Antonelli et al. 2018). Until relatively recently, both long-distance dispersal (Upchurch 2008; Raven and Axelrod 1974; Nelson 1978) and niche evolution among close relatives (Holt and Gaines 1992; Peterson, Soberón J, and Sanchez-Cordero 1999) were thought to be rare; however, empirical studies increasingly demonstrate an important role for these processes across many clade-focused studies (Nathan 2006; Donoghue and Edwards 2014; Yoder and Nowak 2006; Sanmartín and Ronquist 2004; de Queiroz 2005).

The Neotropics are an important region for understanding the joint roles of dispersal and niche evolution in plant biogeography due to its high species richness, extreme habitat heterogeneity, and complex geologic history. The recent geology of the northern Neotropics is marked punctuated by periods of mountain uplift in the Andes (Hoorn et al. 2010; Garzione et al. 2008) and the closing of the Isthmus of Panama (Bacon et al. 2015; O’Dea et al. 2016). These two major events play complementary roles in Neotropical plant biogeography (Antonelli et al. 2009; Gentry 1982). The formation of the Isthmus of Panama represented a continually diminishing barrier to dispersal. Before its emergence, the marine region between Central and South America was difficult to traverse for dispersal-limited terrestrial species, while its completion facilitated the Great American Biotic Interchange, a period marked by high rates of dispersal between continents across a large swathe of tropical rainforest (Bacon et al. 2015). Contrastingly, the Andes, which began their rise in the Paleocene with more recent bursts of mountain building (i.e., 4 to 12 My ago; Garzione et al. 2008), represented an increasingly steep barrier to dispersal throughout their formation, particularly for lineages adapted to the wet, lowland tropical conditions that had previously predominated in the region. Andean uplift changed local topography, continental-scale climate patterns, and landscape configuration (Hoorn et al. 2010). These landscape-scale changes influenced diversification and biogeography in Neotropical plants, whether in montane Andean habitats (Lagomarsino et al. 2016; Särkinen, Pennington, et al. 2012; Pennington et al. 2010; Hoorn et al. 2019) or in extra-Andean habitats, including the Amazon basin (Antonelli et al. 2009; Dick et al. 2012).

Focused clade-based approaches are key in assembling the big picture of Neotropical plant biogeography. Applying comparative methods to well-resolved phylogenies allows researchers to establish biogeographic patterns that may not be discernible at the community level (Cavender-Bares 2019) or within very large clades (e.g., all angiosperms; Donoghue and Edwards 2019). Further, the accumulation of evidence across these studies allows us to make statements about generality— or idiosyncrasy— of biogeographic processes within and between regions (Hughes, Pennington, and Antonelli 2012). We adopt such a clade-based approach to understanding the distribution of *Otoba* (Myristicaceae; Fig. 1), a nutmeg relative that occurs throughout the humid northern Neotropics. *Otoba*’s twelve species are distributed from Nicaragua to Brazil, with the highest species richness in the Chocó and western Amazonia (Santamaría-Aguilar, Jiménez, and Aguilar 2019). They grow in many mesic habitats, including lowland tropical rainforest (both terra firme and floodplain), premontane forest, and cloud forest, spanning a broad elevational range (i.e., 0–2500 m) that includes the highest elevation occurrence in Myristicaceae (Jaramillo-Vivanco and Balslev 2020). Individual species of *Otoba* are among the most common in many rainforests, including *O. parvifolia* in Madre de Díos, Peru (Swamy 2017; Pitman, Vargas, and Terborgh 2017) and Madidí, Bolivia (Macía 2008) and *O. glycycarpa* in Yasuní, Ecuador (Guevara Andino et al. 2017) and high várzea forest of the Amazonian floodplain in Brazil and Bolivia (Wittmann et al. 2006). *Otoba* is one of the ten most abundant genera in western Amazonia (ter Steege et al. 2006; Guevara Andino et al. 2017).

**Fig. 1.**
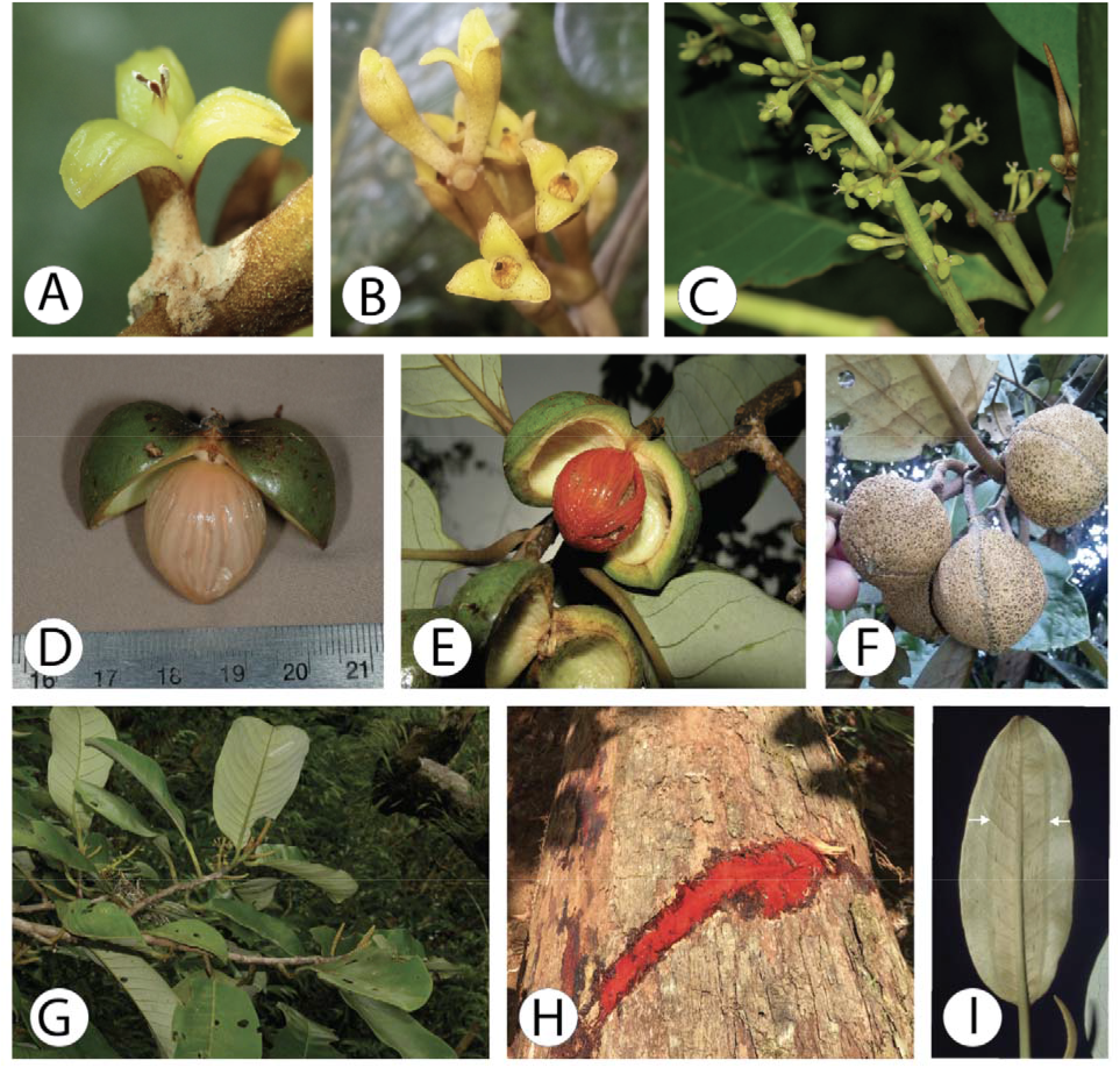
Morphological diversity of *Otoba*. A-C) Floral diversity. A) Staminate and B) pistillate flowers of *O*. *gordoniifolia*; C) Infloresence of Central American *O. novogranatensis*. D-F) Fruit diversity. D) Fruit from South American *O. novogranatensis* showing whitish aril and E) from Central American *O. novogranatensis* showing red aril. F) Unopened capsules of *O. gordoniifolia*. G-I) Vegetative diversity. G) Branch and H) stem cut of Central American *O. novogranatensis*, the latter showing characteristic red exudate. I) Leaf of *O. parvifolia*, showing vernation lines. (Photo credits: A, B, and F by Rudy Gelis, downloaded from iNaturalist with permission; C, E, G, and H by Reinaldo Aguilar; D by Timothy Paine; and I by John Janovec.)

*Otoba*’s broad ecological tolerance and presence across both the Andes and the Isthmus of Panama allow us to use it as a system to understand the roles of dispersal and niche evolution in the establishment of a widespread, ecologically important group. Due to potential dispersal limitation owing to *Otoba*’s large seeds, we hypothesize that niche evolution will be more common than dispersal to disjunct regions to which ancestral lineages are pre-adapted. However, the lack of both a species-level phylogeny of *Otoba* and a robust family-level phylogenetic framework of Myristicaceae impedes placing it into a broader macroevolutionary context. We first perform a biogeographic analysis along the most densely sampled phylogeny of Myristicaceae to date, inferred from publicly available sequence data, to document the family’s movement into the New World. Subsequently, in one of the first target sequence captured-based phylogenomic analyses that relies exclusively on DNA from herbarium specimens collected in the wet tropics, we resolve the phylogeny of *Otoba*. Finally, we use phylogenetic comparative methods to determine the joint roles of ecological niche evolution and dispersal ability across major features of the Neotropical landscape in structuring the biogeographic history of this ecologically important group.

## MATERIALS AND METHODS

### Study clade

Myristicaceae, comprising 21 genera and *ca.* 500 species, is a pantropical plant family with notable importance in ethnobotany, including as food plants (e.g., nutmeg and mace, *Myristica fragrans*), timber species (e.g., *Virola surinamensis*), and hallucinogens (e.g., epená, *Virola* spp.; Alrashedy and Molina 2016). The family is characterized by a strong aromatic scent from essential oils, a pagoda-like growth form (Hallé, Tomlinson, and Zimmermann 1978), dioeciousness with small, trimerous flowers (Armstrong and Tucker 1986) (Fig. 1A-C), red, dilute latex (Fig. 1H), and a characteristic valvate capsule that opens to reveal a large, arillate seed (Fig. 1D-E). These seeds are important food sources for large-bodied birds, primates, and bats, which, in turn, act as seed dispersers (Russo 2003; Forget et al. 2000; Moreira, Riba-Hernández, and Lobo 2017; Giraldo et al. 2007; Melo et al. 2009). Though pollination is less studied, the small, usually imperfect flowers are visited by various small insects including beetles, flies, and thrips in *Myristica* (Sharma and Armstrong 2013; Armstrong and Irvine 1989) and *Virola* (Jardim and Mota 2007); similarly small, generalist pollinators are likely common throughout the family. Further, their abundance in lowland forests (ter Steege et al. 2006) has established Neotropical Myristicaceae an important system for understanding ecological processes that allow species coexistence in hyperdiverse communities in the western Amazon (Queensborough et al. 2007a, b 2007). Within Myristicaceae, *Otoba* is notable for its low-montane distribution, leaves that are folded lengthwise in bud (Fig. 1I), and seeds that most commonly have white arils, not the more typical bright red (as in mace). *Otoba*’s pollen has a unique set of characters that are more similar to African members of the family than other Neotropical genera (Sauquet and Le Thomas 2003).

There has been limited molecular phylogenetic research into Myristicaceae, and no previous research focused on *Otoba*. While many studies support Myristicaceae’s sister relationship to the rest of Magnoliales (Massoni, Forest, and Sauquet 2014; Soltis, Gitzendanner, and Soltis 2007; Qiu et al. 2006), this placement is not consistent across all analyses (Magallón et al. 2015). Three major extant clades are recognized within the family: the malouchoids, pycnanthoids, and myristicoids (Doyle et al. 2004). While most Neotropical members of this family are included in the myristicoids, *Otoba* is currently part of the otherwise African pycnanthoid clade. Phylogenetic relationships within Myristicaceae remain relatively poorly resolved, however, and past studies included very limited taxon sampling (Doyle et al. 2004; Massoni, Couvreur, and Sauquet 2015). Despite a Late Cretaceous origin of stem Myristicaceae (Magallón et al. 2015), molecular dating analyses point to a relatively recent origin of the extant members of the family in the Oligocene (Massoni, Couvreur, and Sauquet 2015). Fossil evidence, in the form of the seed *Myristicacarpum chandlerae* in the London Clay formation that displays characteristics of crown Myristicaceae, suggests that the extant radiation of the family has existed since at least the early Eocene (Doyle, Manchester, and Sauquet 2008).

### Taxon Sampling

Twenty accessions of Otoba representing eleven of the twelve accepted species (i.e., O. acuminata, O. cyclobasis, O. glycycarpa, O. gordoniifolia, O. gracilipes, O. latialata, O. novogranatensis, O. parvifolia, O. scottmorii, O. squamosa, and O. vespertilio) were sampled. All accessions came from herbarium specimens; voucher information is available in Appendix S1 (see Supporting Information). Herbarium acronyms follow Index Herbariorum (Thiers, constantly updated: http://sweetgum.nybg.org/science/ih/). To serve as outgroups, data from the following transcriptomes available on 1KP project (Carpenter et al. 2019; One Thousand Plant Transcriptomes Initiative 2019); <https://db.cngb.org/onekp/>) were gathered for Myristicaceae (Myristica fragrans), the broader Magnoniales (Magnolia maudiae, Annona muricata), and Laurales (Cassytha filiformis, Sassafras albidum, and Persea borbonia).

To put *Otoba* in a broader phylogenetic context, we also inferred a phylogeny of Myristicaceae using publicly available DNA sequences. For this, we sampled plastid loci matK, rbcL, and ndhF from 53 species of Myristicaceae represented in GenBank, as well as outgroup Annonaceae, Magnoliaceae, Degeneriaceae, and Lauraceae (Appendix S2).

### DNA extraction, library preparation, target enrichment, and sequencing

500 mg of leaf tissue was homogenized using an MP Biomedicals FastPrep-24TM 5G. DNA extraction followed a modified sorbitol extraction protocol (Štorchová et al. 2000). DNA concentration was quantified using an Invitrogen Qubit 4 Fluorometer, and fragment size was assessed on a 1% agarose gel. For samples with a high concentration of large fragments (>800 bp), DNA was sheared using a Bioruptor Pico (Diagenode Inc., Denville, New Jersey, USA) to obtain an average fragment size of ~500 bp. Library preparation was carried out using KAPA Hyper Prep and HiFi HS Library Amplification kits (F. Hoffmann-La Roche AG, Basel, Switzerland) and with iTru i5 and i7 dual-indexing primers (BadDNA, University of Georgia, Athens, Georgia, USA). Library preparation with KAPA Hyper Prep followed the manufacturer’s protocol (KR0961 – v8.20) with the following modifications: reaction volumes were halved (i.e., 25 μL starting reaction) and bead-based clean-ups were performed at 3X volume rather than 1X volume to preserve more small fragments from degraded samples. Library amplification reactions were performed at 50 μL. Target enrichment was carried out using the MyBaits Angiosperms353 universal probe set (Däicel Arbor Biosciences, Ann Arbor, MI; Johnson et al. 2019) following the protocol outlined in Hale et al. (2020). Twenty nanograms of unenriched DNA library was added to the cleaned, target enriched pool to increase off-target, chloroplast fragments in the sequencing library. DNA libraries were sent to Novogene (Sacramento, California, United States) for sequencing on an Illumina HiSeq 3000 platform with 150 bp paired-end reads.

### Sequence processing, assembly, and alignment

Raw sequence reads were demultiplexed by Novogene. Adapter sequence removal and read trimming were performed using illumiprocessor v2.0.9 (Faircloth et al. 2012; Faircloth 2016), a wrapper for trimmomatic v0.39 (Bolger, Lohse, and Usadel 2014). The default settings were used and reads with a minimum length of 40 bp were kept.

HybPiper v1.3.1 (Johnson et al. 2016) was used to assemble and extract target regions. Read mapping, contig assembly and coding sequence extraction were performed running the reads_first.py script. The intronerate.py script extracted introns and intergenic sequences flanking targeted exons. The retrieve_sequences.py script was run first with the “dna” argument to extract coding regions and subsequently with the “supercontig” argument to extract both coding and non-coding regions as a single concatenated sequence for each target gene. Individual genes were aligned using default parameters in MAFFT v7.310 (Katoh and Standley 2013). Alignments were visually inspected in AliView v1.18.1 (Larsson 2014). Obvious alignment errors were manually corrected and assembly errors (identified by high divergence on terminal ends of contigs), as well as areas with substantial interspecific variation that were difficult to align, were removed from individual alignments. Outgroup sequences were added to cleaned alignments and aligned using default parameters MUSCLE v3.8.31 (Edgar 2004) in AliView (Larsson 2014). Summary statistics on gene alignments were obtained using AMAS (Borowiec 2016), including length, missing data, and number of parsimony informative sites.

Off-target chloroplast reads were extracted using FastPlast v1.2.6 (https://zenodo.org/record/973887). For all samples there was insufficient data to produce a full plastome. The SPAdes-assembler built into FastPlast iteratively used k-mer lengths of 55, 87, and 121. Assembled contigs from the 87 k-mer length iteration were mapped to a reference plastome obtained from GenBank (Clark et al. 2016: *Horsfieldia pandurifolia,* GenBank accession number NC_042225.1). Series of incongruous basepairs at the ends of contigs were considered to be the result of assembly error and were cleaned by eye before generating a consensus sequence. Consensus sequences for each sample were aligned visually against the *Horsfieldia* plastome. Plastomes of *Annona muricata* (MT742546.1), *Cassytha filiformis* (MF592986.1), *Magnolia maudiae* (MN990580.1), and an unverified plastome for *Myristica yunnanensis* (MK285565.1) were added as additional outgroups.

Plastid sequences from Myristicaceae from GenBank for whole-family phylogenetic analysis were aligned using default parameters in MUSCLE in AliView.

### Phylogenetic analysis and divergence time estimation

#### Otoba

Alignments for each Angiosperm353 locus and the plastome were processed with trimAl (Capella-Gutiérrez and Silla-Martínez 2009), assigning a gap threshold of 15% or 20% to each column, depending on the number of taxa in the alignment. Thresholds were chosen to maintain columns with data for four or more total accessions. A concatenated alignment of all nuclear and plastid data was analyzed in RAxML under the GTR model with optimization of substitution rates and site-specific evolutionary rates. RogueNaRok v1.0 (Aberer, Krompaß, and Stamatakis 2011) was subsequently used to identify individuals that negatively impacted phylogenetic inference. Individuals identified by RogueNaRok and those with little data (total bp <1% of aligned length) were excluded from final analysis. Coalescent-based phylogenetic methods were not attempted due to the small number of loci that were successfully recovered for multiple species, as well as differences in success in capture plastid sequences across taxa.

Divergence times were estimated on the inferred topology using penalized likelihood in treepl (Smith and O’Meara 2012). Confidence intervals were calculated along 100 bootstrap replicates from a RAxML analysis and summarized using TreeAnnotator (Bouckaert et al. 2019). The large size of the *Otoba* phylogenomic dataset precluded a full Bayesian analysis, as performed for Myristicaceae (see below). Crown ages from the literature (Magallón et al. 2015; Massoni, Couvreur, and Sauquet 2015) for Laurales + Magnoliales, Laurales, Magnoliales, and Myristicaceae were applied as secondary calibrations. Because Massoni, Couvreur, and Sauquet (2015) presented five different calibration schemes, we used the youngest and oldest dates in the 95% confidence interval across their different analyses as the minimum and maximum ages in our analysis. We performed an additional set of analyses that included a secondary calibration on the crown node of *Otoba* that corresponded to the 95% highest posterior density (HPD) of this node in the more densely sampled phylogeny of the entire Myristicaceae, which used calibrations derived from the same two sources (see below). Calibration schemes for all dating analyses can be found in Appendix S3.

#### Myristicaceae

We performed preliminary ML analyses on individual loci in RAxML. Sequences that systematically reduced support values were identified in RougeNaRok (i.e., RNR values >0.5) and removed. Updated alignments were then concatenated and used to jointly infer species relationships and divergence times in BEAST v2.3.6 (Bouckaert et al. 2019). We used PartitionFinder2 (Lanfear et al. 2017) to identify an appropriate partitioning scheme and model of molecular evolution for each partition. For each of two analyses, we assigned normally distributed priors nodes corresponding to the same calibration points used for dating in *Otoba*, with calibration points from both Magallón et al. (2015) and Massoni, Couvreur, and Sauquet (2015). For each analysis, we linked tree and clock models and set a lognormal clock prior. Eight runs each with four chains were allowed to progress for 10 million generations, sampling every 5,000 generations. Convergence was assessed using ESS values in Tracer v1.7.1 (Rambaut et al. 2018) with a cutoff value of 200. Maximum clade credibility trees were assembled in TreeAnnotator on the combined output from all runs after a 20% burnin was discarded.

### Biogeographic inference

We inferred biogeographic history of *Otoba* using BioGeoBEARS (Massana et al. 2015; Matzke 2014) along the phylogeny that was time-calibrated using ages from Magallón et al. (2015) and the crown node of *Otoba* from our Myristicaceae-wide analysis. Movement between both 1) Central and South America and 2) across the Andes were modeled. Each species was coded for 1) occurrence in Central America, South America, or both, as well as for 2) their distribution on the western side of Andes, including the Darién gap and Central America, or the eastern side of the Andes including western Amazonia. Six biogeographic models (i.e., DIVA-like, BayAREAlike, and DEC, as well as the addition of jump dispersal for each of those models) were tested; using AICc scores, the DIVA-like model was selected for both reconstruction of continental movements and reconstruction of distribution around the Andes. A maximum of two ancestral areas was allowed for both analyses.

To put *Otoba* in a global context, we performed a biogeographic analysis along the phylogeny of Myristicaceae calibrated with dates from Magallón et al. (2015). Outgroups, which had non-representative sampling, were removed. The range of each species was coded as Asia, Africa, or the Neotropics. Analyses were performed as above, with the DIVA-like model selected as best-fit.

### Environmental niche modeling and evolution

Environmental niche models were generated for each species. Occurrence data were gathered from GBIF (https://gbif.org). Using the R package raster (Hijmans et al. 2015), the standard WorldClim 2.0 30s Bioclimatic variable layers (Fick and Hijmans 2017) and the WorldClim 2.1 30s elevation layer (https://www.worldclim.org/data/worldclim21.html) were stacked and clipped to the tropical latitudes of the Americas (extent = −120, −30, −23, 23). Variables were assessed for correlation using R package ENMTools (Warren et al. 2019). Bioclimatic variables for diurnal range (BIO 2), isothermality (BIO 3), maximum temperature of warmest month (BIO 5), precipitation seasonality (BIO 15), precipitation of warmest quarter (BIO 18), and precipitation of coldest quarter (BIO 19) were chosen as minimally correlated variables for modeling. Generalized linear models were prepared for each species with ENMTools. Because phylogenetic results suggested that *O. novogranatensis* is not monophyletic, occurrences from Central America and South America were analyzed separately.

To quantify the multidimensional niche in *Otoba*, a phylogenetic principal components analysis of the average value of the 19 WorldClim bioclim variables using a correlation matrix was performed using the phyl.pca function in phytools (Revell 2012). Ancestral state reconstruction of the first three principal components was performed with the contMap function in phytools (Revell 2012). We then used Orstein-Uhlenbeck models in the l1ou R package (Khabbazian et al. 2016) to detect shifts to different climatic niches in the absence of an *a priori* hypothesis about their location or convergent regimes. We searched for shifts in the first three climate principal components on the *Otoba* phylogeny, and assessed support for the shifts identified via a bootstrap analysis with 100 replicates.

## RESULTS

### Summary statistics of data assembly

The number of Angiosperm353 loci captured, average sequence length, number of ungapped basepairs of plastome for each sample, and collection year are listed in Appendix S4. A heatmap of the percent of the reference protein length recovered for each sample at each locus can be found in Appendix S5. Due to large amounts of missing data in both nuclear and chloroplast regions, the following samples were excluded from all phylogenetic analyses: *O. gracilipes*_DC884, *O. novogranatensis*_EB500, *O. parvifolia*_DN9151, *O. vespertilio,* and two recently described species (i.e., *O. sp. nov.* RC5752 [= *O. scottmorii*] and *O. sp. nov.*_JP16902 [= *O. squamosa*]; Santamaría-Aguilar and Lagomarsino 2021). Overall, we were able to include 7 of the 12 described species of *Otoba*, representing 13 accessions, in phylogenetic analyses.

Hybrid enriched target sequence capture success was variable across samples. Half of the 20 samples submitted for sequencing recovered fewer than 10 Angiosperm353 loci; only 3 samples recovered more than 100 loci (Appendix S4). Success in gathering off-target plastid data did not necessarily correspond to success in capturing nuclear loci. For example, the sample with the most nuclear data– *O. novogranatensis*_WS36336 with 217 of the 353 targeted loci– did not recover useful plastome data. On the other hand, nearly half of the chloroplast genome was obtained for *O. parvifolia*_MS1182, despite recovering only 6 nuclear loci.

### Phylogenetic analyses of Otoba

We found support for the monophyly of *Otoba* within Myristicaceae (Bootstrap support [BP]= 100; Fig. 2) and for three subclades of *Otoba* (Fig. 2). The first subclade (BP=88), corresponding to South American accessions of *O. novogranatensis*, is sister to the rest of *Otoba* (BP =100). The second subclade includes a paraphyletic *O*. *parvifolia* and a single accession of *O*. *glycycarpa* (BP= 75), while the third comprises *O. acuminata*, *O. cylclobasis, O. gordoniifolia*, *O. latialata*, and *O. novogranatensis* from Central America (BP= 100). Within the third subclade, we find that *O. cyclobasis* is sister to the remaining species, with *O. gordoniifolia* and *O. latialata* forming a sister pair and *O. acuminata* sister to Central American accessions of O. novogranatensis.

**Fig. 2.**
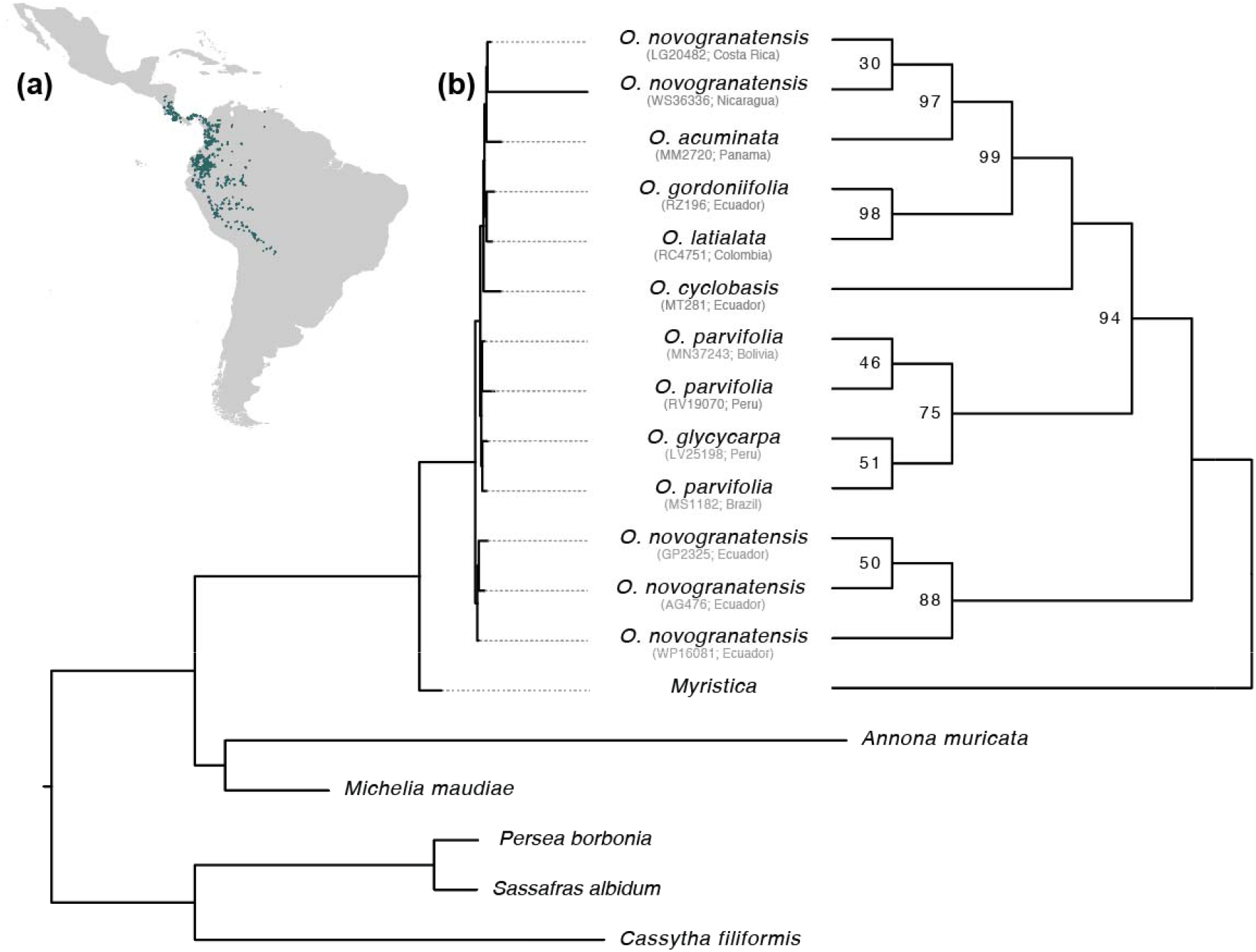
Phylogeny of Otoba and its distribution within the Neotropics. (a) Distribution map of Otoba made using points from GBIF. (b) Results of a maximum likelihood phylogenetic analysis of concatenated chloroplast and nuclear data in RAxML. Both topologies are identical; the tree at left shows branch lengths, while the cladogram on the right shows more clearly depicts the branching relations in the ingroup and includes support values at nodes with <100 bootstrap support. Nodes that lack support values had maximal support. Collection number and country are noted below each ingroup taxon name.

Divergence times in *Otoba* estimated from different calibrations schemes were similar between calibrations from Massoni et al. (2015) and Magallón et al. (2015), but differed if an additional calibration derived from our BEAST analyses of Myristicaceae were used (Appendix S6). All estimates support the radiation of *Otoba* in the Miocene (Appendix S6). The crown age for *Otoba* based only on calibrations from Magallón et al. (2015) is estimated to be 7.32 Ma [3.2–49.1] and based only on Massoni, Couvreur, and Sauquet (2015) is 8.23 Ma [4.3–73.6]. These dates were pushed back when the additional calibration was added, to 12.48 Ma [3.8– 34.8] with the Magallón et al. (2015) calibrations and 13.74 Ma [4.3–30.5] with those from Massoni et al. (2015).

### Phylogenetic analyses of Myristicaceae

The monophyly of Myristicaceae is well-supported (PP=1.0 with Massoni calibrations/1.0 with Magallón calibrations), as is the monophyly of four major subclades that correspond to broad geographic regions (Fig. 3; Appendices S7-8). An African (including Malagasy) subclade (PP=1.0/1.0), comprising the genera *Pycnanthus*, *Coelocaryon, Staudia*, *Brochoneura*, *Maulotchia*, and *Cephalosphaera*, is sister to the rest of the family. The first of two Asian subclades (PP= 0.64/0.66) comprises the genus *Horsfieldia* and is sister to the second Asian subclade + the Neotropical subclade. The second Asian subclade (PP=0.84/0.84) includes three genera, each found to be monophyletic in our sampling: *Knema* (PP=1.0/1.0), *Myristica* (PP=1.0/1.0), and *Gymnacanthera* (PP=1.0/1.0). Finally, the Neotropical subclade (PP=0.99/0.99) includes all Neotropical genera. Within it, *Otoba* (PP=1.0/1.0) and *Virola* (PP=1.0/1.0) are subsequently sister to the remaining genera, with *Iryanthera* (PP=1.0/1.0) forming a clade sister to a poorly supported *Composoneura* + *Osteophloeum* (PP=0.47/0.45).

**Fig. 3.**
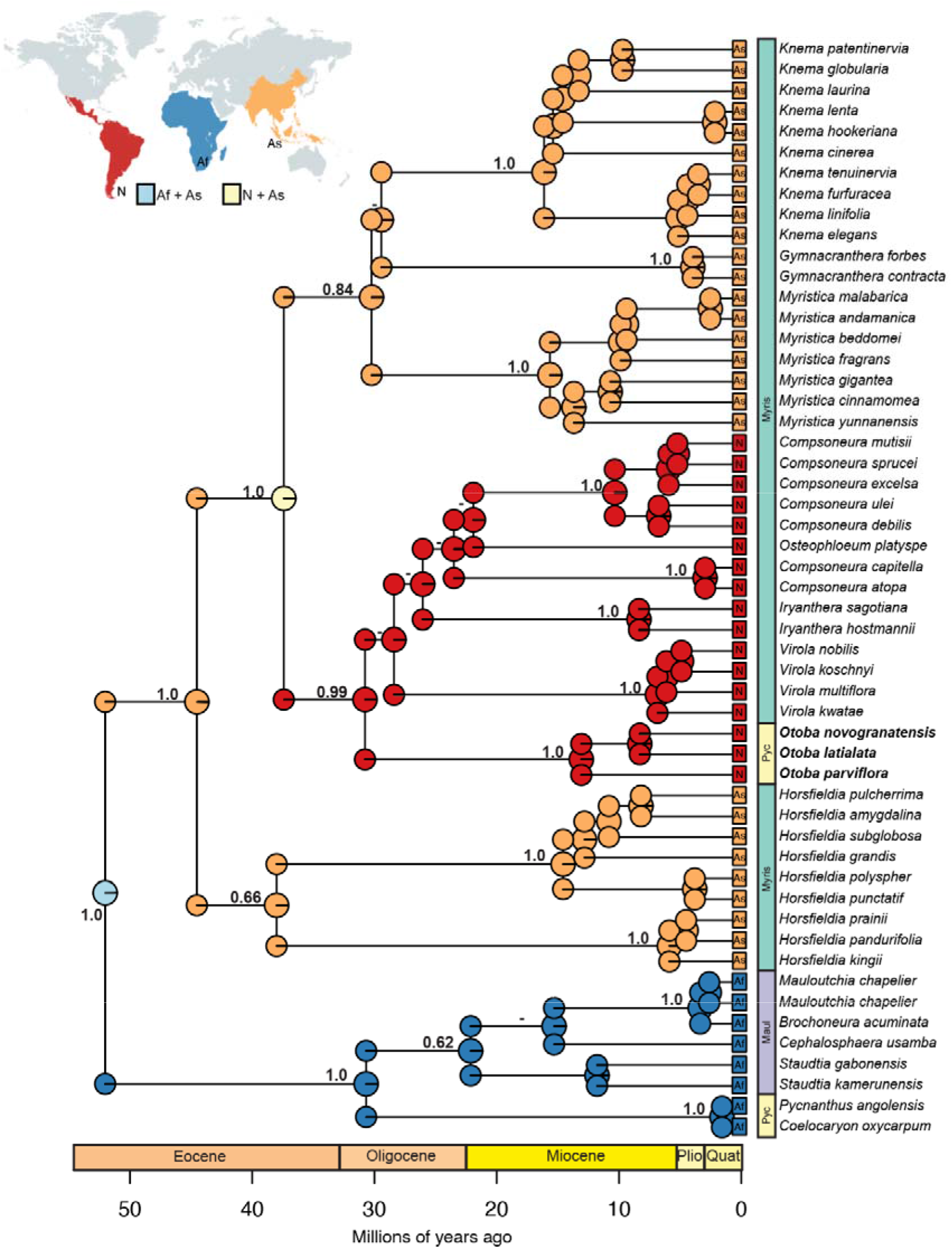
Time-calibrated plastid phylogeny of Myristicaceae depicting global biogeography. The phylogeny represents the maximum clade credibility tree of a BEAST2 analysis using calibration points from Magallón et al (2015). Values at nodes represent posterior probabilities, with dashes representing <50 PP; support for relationships below genus level are not included except in instances of deep splits with a genus, but are available in Appendix S7, as are the 95% HPD for the date at each node. Ancestral area reconstructions as inferred by the best-fit DIVA-like model in BioGeoBEARS are depicted in pie charts at each node, with colors corresponding to the areas in the map at top left (dark blue= Africa; orange = Asia; red= Neotropics; yellow = both Neotropics and Asia; light blue = both Africa and Asia). The previously understood major clades of Myristicaceae from Sauquet et al. (2003) are indicated by the colored bars at the right of the phylogeny (yellow = pycnanthoids; purple = mauloutchioids; green = myristicoids). Color coding of the geologic timescale follows the Commission for the Geological Map of the World (abbreviations: Quat = Quaternary; Plio= Pliocene)

As within *Otoba*, we find that the Massoni et al. (2015) calibration scheme gives slightly younger ages (Appendices S7-8). Crown Myristicaceae is estimated to have originated in the Eocene in both analyses, at 40.77 Ma [26.9–54.7] with the Massoni et al. (2015) calibrations and at 52.02 Ma [26.25–78.91] with the Magallón et al. calibrations. The crown Neotropical subclade originated in the Paleocene in both analyses (26.45 Ma [16.28–38.27] / 30.77 Ma [16.77–49.07]). Ages inferred for *Otoba* in the Myristicaceae-wide dataset are older than in the *Otoba*-specific analyses (Appendix S6), but still within the Miocene (11.41 Ma [2.81–21.62] vs.13.08 Ma [2.18–25.74]).

### Biogeographic Reconstruction

We find that Myristicaceae, as a whole, originated in the Old World tropics, with a broad range spanning both the Asian and African tropics. It expanded into the Neotropics once, between 37.41 and 30.77 Ma via an ancestor with a broad range that included the Americas and Asia (Fig. 3). Subsequent *in situ* diversification resulted in five endemic extant Neotropical genera, including *Otoba*. A South American ancestor (Fig. 4A) is inferred for *Otoba*, with expansion into Central America via widespread ancestors. A single dispersal from the western to the eastern side of the Andes explains the divergence of the *O. glycycarpa*-*O. parvifolia* clade from the rest of the genus (Fig. 4B).

**Fig. 4.**
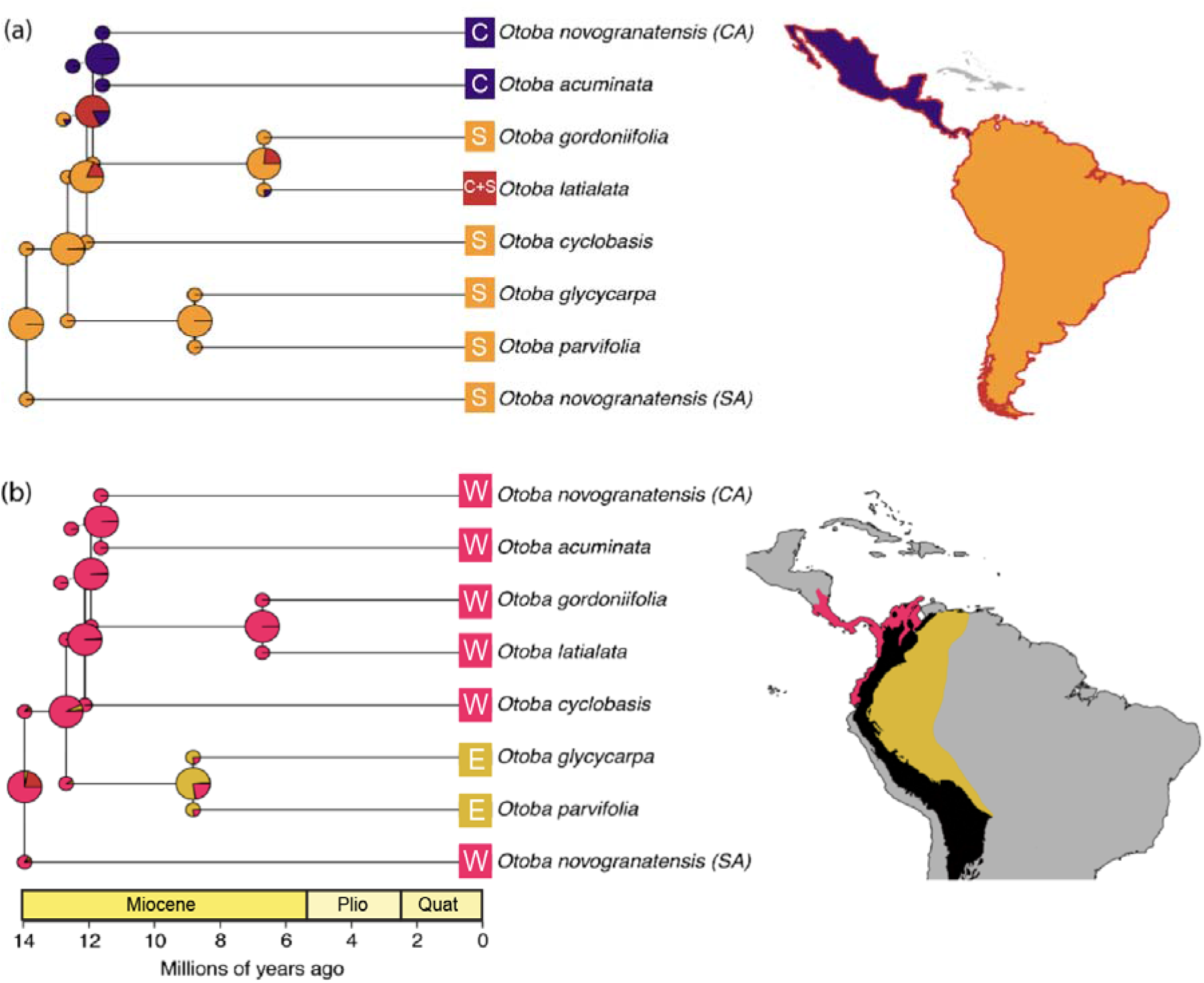
Ancestral area reconstructions in *Otoba* for (A) Central America versus South America and (B) western Andes and Central America versus eastern Andes and the Amazon. Pie charts at nodes display the probability that the ancestor occupied a given range. Maps to the left of each tree are color coded to correspond with the geographic areas coded on the tree. In (A), orange = South America, purple = Central America, and red = both Central and South America; the distribution of each species in Central and/or South America is additionally reflected in the color-coded boxes at the tips of the tree. In (B), pink = western Andes and Central America, and mustard = eastern Andes and the Amazon; the Andes are represented in black. Color coding of the geologic timescale follows the Commission for the Geological Map of the World (abbreviations: Quat = Quaternary; Plio= Pliocene).

### Niche Modeling and Evolution

Visualizations of GLM models can be found in Fig. 5a and average values for each BIOCLIM variable and elevation in Appendix S9. The first three principal components from our phylogenetic PCA of BIOCLIM variables describe 91.7% of the variation in climatic preferences. The most important loadings for pPC1, which describes 57.9% of the variation, are mean temperature of the driest quarter (BIO9), mean temperature of the coldest quarter (BIO11), mean annual temperature (BIO1), minimum temperature of the coldest month (BIO6), and mean temperature of the warmest quarter (BIO10), making it a useful proxy for temperature regime (Appendix S10). The most important loadings for pPC2, which describes 21.3% of the variation, are temperature seasonality (BIO4) and isothermality (BIO3), making it a useful proxy for temperature stability (Appendix S10). Finally, the most important loadings for pPC3, which describes 12.5% of the variation, are mean diurnal temperature range (BIO2), annual temperature range (BIO7), and precipitation seasonality (Appendix S10). We observed a broad occupation of climate phylomorphospace (Fig 5c). Of particular note, the sister species *O. latialata* (native to the Chocó region) and *O. gordoniifolia* (an Andean montane species) are the two most distinct species, falling on extreme ends of pPC1 and also differing substantially in pPC2. Further, the western Amazonian sister pair *O. glycycarpa* and *O. parvifolia*, while relatively similar to each other, form a separate cluster in phylomorphospace, largely due to their extreme value of pPC3.

**Fig. 5.**
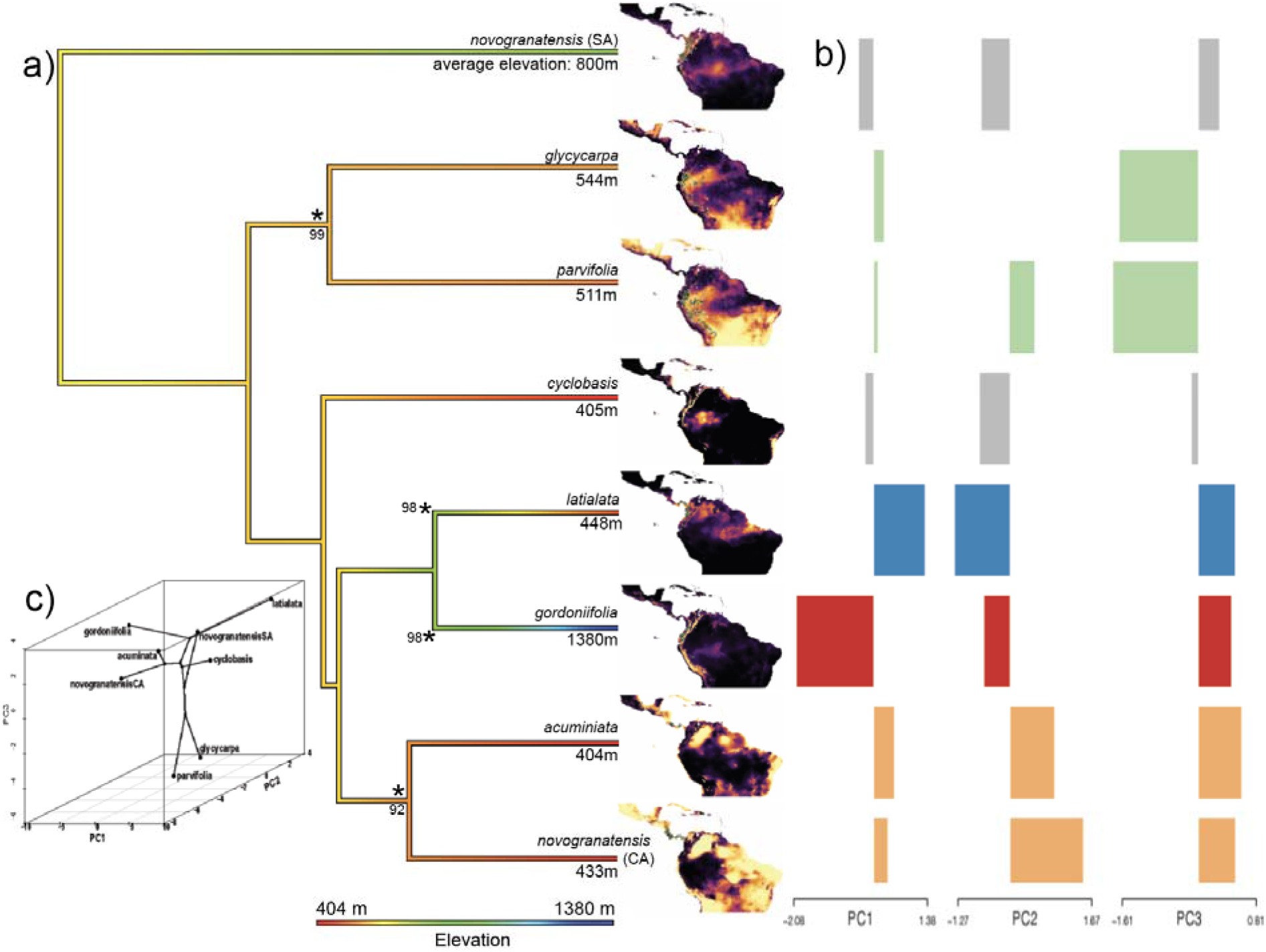
Niche evolution in *Otoba* is dynamic. A) Ancestral state reconstruction of average elevation (m) in *Otoba* shows multiple shifts in elevational range from low montane ancestor, with two shifts to increased elevations and at least one shift towards lower elevations. Colors at tips demonstrate the wide range of average elevational preferences among species, ranging from 404m in *O. acuminata* to 1380m in *O. gordoniifolia*. At right, species distributions modeled in Maxent demonstrate the variation in potential niche suitability in the Neotropics. B) Orstein-Uhlenbeck models implement in l1ou infer four shifts in climate space, denoted by asterisks at nodes with bootstrap values, corresponding to the eastern Amazonian clade (i.e., *O. glycycarpa* and *O. parvifolia*); *O. latialata*; *O. gordoniifolia*; and a Central American clade comprising *O. acuminata* and *O. novogranatensis* (CA). C) Phylomorphospace of phylogenetic PCA of BIOCLIM variables shows broad occupation of climate space, including multiple instances where close relatives differ along at least one axis

Orstein-Uhlenbeck models in l1ou identify four highly supported niche shifts in our analysis of pPC1-3 (Fig. 5b). These correspond to 1) *O. glycycarpa* + *O. parvifolia* (PP=99), which experienced a shift to increased temperature extremes and decreased precipitation seasonality in pPC3; 2) *O. latialata* (PP=98), which experienced a shift to warmer temperatures (pPC1) and increased temperature stability (pPC2); *O. gordoniifolia* (PP=98), which experienced a shift towards cooler temperatures (pPC1); and *O. acuminata* + *O. novogranatensis* (PP=92), which experienced a minor increase in temperature (pPC1) and a larger shift towards decreased temperature stability (pPC2). These shifts mirror results from ancestral state reconstruction of individual pPCs (Appendix S11), and are often accompanied by shifts in biogeography (Fig. 4,5).

## DISCUSSION

### Phylogeny of Otoba and the broader Myristicaceae

Our plastid phylogeny of Myristicaceae, which combined publicly available plastid sequences with the same loci from our target capture data, is the most densely sampled phylogeny of the family to date (Fig. 3). At the deepest temporal scale, our results are broadly similar to past studies (Sauquet et al. 2003; Doyle et al. 2004): we resolve an African subclade that includes the mauloutchoids and African pycnanthoids as sister to the Asian and American myristicoid clade. However, there are some key differences. Notably, we find that *Otoba* is part of the Neotropical subclade of myristicoids, and not sister to the African pycnanthoid genera as in Sauquet et al. (2003). This result has important biogeographic implications, discussed below, and suggests that informal clade names previously applied to Myristicaceae need redefinition. Further, resolution within the myristicoid clade is improved (Fig. 3). We find support for three subclades: 1) the Asian genus *Horsfieldia*; 2) the remaining Asian genera (i.e., *Gymnacanthera*, *Knema*, and *Myristica*); and 3) all Neotropical genera (i.e., *Compsoneura, Iryanthera*, *Osteophloem*, *Otoba*, and *Virola*). This updated phylogeny provides a useful framework for future macroevolutionary research in Myristicaceae.

While our family-wide analysis found that *Otoba* is sister to the remaining Neotropical genera, our species-level phylogenomic analysis resolved relationships within the genus. This represents the first phylogeny of *Otoba* to date. Phylogenetic results within *Otoba* point to the need for future phylogeographic and taxonomic study: there is strong evidence for polyphyly of the widespread *O. novogranatensis,* with two distinct lineages corresponding to South American and Central American accessions. While there was no reason to expect that *O. novogranatensis* represented more than a single species from previously published taxonomic research (Jaramillo-Vivanco and Balslev 2020; Jaramillo, Muriel, and Balslev 2004; de Candolle 1856), our results demonstrate that it represents at least two cryptic species with disjunct geographic distributions. A preliminary revision of herbarium specimens reveal that South American individuals morphologically differ from Central American individuals in their thicker pericarp, pubescent ovaries, and generally white arils.

### Challenges of herbarium phylogenomics from wet tropical specimens

As it is often not feasible (or safe) to collect living material for the taxonomic and/or geographic breadth of many clades, reliance on DNA from herbarium specimens is often necessary to achieve dense taxonomic sampling in phylogenetic studies of widespread tropical groups. However, even high efficiency target capture will fail when DNA is low quality, as is typical in herbarium specimens collected in wet tropical ecosystems (Brewer et al. 2019). Molecular research is made even more difficult when specimens are preserved in ethanol ahead of drying (Särkinen, Staats, et al. 2012)— a common scenario for *Otoba*, and likely universal in the herbarium specimens that we sampled. It is thus not surprising that not all of our samples generated useful sequences. To extend the potential utility of newly collected specimens for future genomic research, efforts should be made to collect leaf tissue in silica gel or apply another preservation technique (Funk et al. 2017).

The Angiosperms353 probeset is proving to be an exciting toolkit that plant systematists are using to successfully resolve relationships across taxonomic groups and phylogenetic depth (Slimp et al. 2021; Johnson et al. 2019), reminiscent of ultraconserved elements in animals (Faircloth et al. 2012). However, universal probesets result in different quality datasets depending on phylogenetic scale and distance from species used in their design (Hutter et al. 2019). While we were able to infer relationships in *Otoba* using Angiosperms353 loci, very limited variation across our dataset (i.e., the proportion of variable sites in concatenated target loci was 0.285) and inconsistent capture, with most accessions recovering a fewer than a third of loci, likely resulted in low support values in our phylogenetic analyses. It is likely that a custom probe kit designed for *Otoba* and close relatives would have outperformed the Angiosperms353 loci, either alone or in combination (Jantzen et al. 2020; Ogutcen et al. 2021), especially as only a single species of Myristicaceae was used in the development of Angiosperms353 loci (Johnson et al. 2019). However, developing such custom loci is predicated on the existence of genomic resources, either pre-existing or newly generated, which would have been difficult for *Otoba*. While the number of angiosperm genomic resources is constantly growing, there are still no transcriptomes or nuclear genomes available for *Otoba* and data is limited for Myristicaceae overall: the only public resource is a single transcriptome in the 1KP database (nutmeg, *Myristica fragrans*). Because this scenario is common— especially in tropical plant groups, which tend to be understudied (Goodwin et al. 2015)— the universal utility and subsequent promise of assembling a standardized set of loci across analyses is desirable.

### Biogeography of Myristicaceae is marked by relatively few major movements

As a whole, Myristicaceae have a straightforward biogeographic history with relatively few movements between regions and potentially no long-distance dispersals. At a continental scale, we infer fewer major biogeographical events than previously postulated (Doyle et al. 2004): one range expansion and two range contractions (Fig. 3). The relatively young age of Myristicaceae precludes a role of a Laurasian ancestor in the formation of its tropical disjunctions, contrasting with many of its relatives in Magnoliales, including Magnoliaceae (Azuma et al. 2001). Instead, we find that from a widespread common ancestor in the eastern hemisphere during the early Eocene, distinct lineages became restricted to Africa and Asia by the mid-Eocene. A subsequent lineage expansion to include the Neotropics occurred during the late Eocene-Oligocene, eventually giving rise to all extant Neotropical Myristicaceae, which are embedded within an otherwise Asian clade.

The Oligocene timing of Myristicaceae’s amphi-Pacific disjunct is consistent with the Boreotropics hypothesis (Wolfe 1975). This is the scenario in which plant lineages with widespread distributions in the warm, wet conditions of the Northern Hemisphere in the Late Paleocene-Eocene were disrupted by climatic cooling in the Oligocene, resulting in climate-driven vicariance (Lavin and Luckow 1993). Fossil evidence is further consistent with a Boreotropical distribution for Myristicaceae. This includes the occurrence of *Myristicacarpum chandlerae*, which shares morphological traits with both Old World and New World taxa, in the early Eocene of the London Clay flora (Doyle, Manchester, and Sauquet 2008), as well as leaves in the middle to late Eocene of the Alaska Gulf (Wolfe 1977). Support for a potential role of range expansion over continental plates in close proximity is further bolstered by lack of long-distance dispersal events in the history of the family, as well as the presence of large seeds for which long-distance dispersal is improbable. However, it is impossible to rule out long-distance dispersal over water without additional fossil evidence. Similar intercontinental, amphi-Pacific tropical disjunctions with documented or likely occurrence in the Boreotropics are seen in Annonaceae (Xue et al. 2018; Thomas, Tang, and Saunders 2017), Melastomataceae (Morley and Dick 2003), Araliaceae (Valcárcel and Wen 2019), and other groups.

Finally, *Otoba* is strongly supported as an *in situ* radiation within a larger Neotropical clade. Contrasting with previous studies within Myristicaceae with more limited taxon sampling (Doyle et al. 2004), our updated phylogenetic and biogeographic results demonstrate that intercontinental long-distance dispersal does not need to be invoked to explain the origin of *Otoba*.

### Landscape change, niche evolution, and dispersal limitation interact to drive biogeography in Otoba

The relatively stable global biogeography of Myristicaceae contrasts with the dynamic history of *Otoba* within the Neotropics. At the time of origin of Neotropical Myristicaceae, a broad distribution across the northern Neotropics would have been facilitated by a more-or-less contiguous swathe of lowland rainforest and the low height of the Andes, including multiple low elevation gaps (Hoorn et al. 2010). However, by the time that crown *Otoba* originated in the late Miocene to early Pliocene, the genus was restricted to the western side of the Andes. (Fig. 4), an ancestral range shared with relatively few other Neotropical tree groups, including a clade of Annonaceae (Pirie et al. 2018). By this time, the northern Andes had gained approximately half their elevation (Garzione et al. 2017) and analogs to modern montane cloud forests had formed (Hughes 2016; Martínez et al. 2020). Thus, even though they had not yet reached their full height, the Andes would have represented a significant barrier to dispersal for low- to-mid-elevation groups like *Otoba.* This is supported in our results: the Andes structure biogeography in *Otoba*, with species and subclades occurring on only one side of the mountain chain. A movement to the eastern slope occurred only once in *Otoba* at *ca*. 9 Ma, resulting in an eastern Andean/Amazonian clade comprising the two widespread species *O. parvifolia* and *O. glycycarpa*. This movement could be explained by either a dispersal across the Andes or a vicariance scenario involving the final uplift of the Mérida cordillera in Venezuela.

Movement between Central and South America is more dynamic than across the Andes in *Otoba.* This is not surprising as the Isthmus of Panamá was either formed (Montes et al. 2012; Bacon et al. 2015) or in the process of forming (O’Dea et al. 2016) when the genus originated, providing a land connection of appropriate habitat type that could facilitate northward movement. From an ancestral range of South America, a range expansion to include Central America is inferred for the most recent common ancestor of *O. acuminata*, *O. novogranatensis* (CA), *O. gordoniifolia*, and *O. latialata* in the late Miocene (Fig. 4). *Otoba acuminata* and Central American *O. novogranatensis* subsequently became restricted to Central America, while *O. latialata* has a widespread distribution in the Chocó-Darién moist forest from Colombia to Panama. It is possible that the common ancestor of this subclade may have inhabited a similar distribution to *O. latialata*. If this is the case, it may represent subset sympatry—when one daughter lineage inherits the ancestral range and the other daughter(s) inherit a portion of the ancestral range (Ree et al., 2005). An additional Central American species, *O. vespertilio*, was not included in our phylogenetic analysis due to poor data quality (Appendix S4), but morphological evidence suggests this species likely represents an independent colonization of Central America (Santamaría-Aguilar, Jiménez, and Aguilar 2019). These migrations occurred within the last 10 million years, a timeframe that supports the role of the closure of the Isthmus of Panama (Montes et al., 2012; Bacon et al., 2013).

The limited movement of *Otoba* is likely explained by poor dispersal ability— a pattern echoed across the phylogeny of Myristicaceae, in which long-distance dispersal played little to no role in its current distribution (Fig 3). *Otoba*’s relatively large seeds, which are dispersed by birds, primates, and bats (Santamaría-Aguilar, Jiménez, and Aguilar 2019; Melo et al. 2009; Nuñez-Iturri and Howe 2007; Giraldo et al. 2007), make dispersal events over water barriers or large stretches of unsuitable terrestrial habitat uncommon compared to groups that are dispersed by wind (Pérez-Escobar et al. 2017) or migratory passerine birds (Nathan et al. 2008). Further, dispersal ability in tropical trees is known to be negatively impacted by seed size (Muller-Landau et al. 2008). On a continental scale and over evolutionary time, this has resulted in remarkably few long distance biogeographic movements. *Otoba*’s migration into Central America was likely facilitated by the continuous land bridge of suitable habitat following the closure of the Isthmus of Panama, while cold high-elevation habitats likely prevented more frequent traversing of the Andes. Further supporting the role of low dispersal ability in the biogeography of *Otoba*, the genus does not occupy all of the suitable habitat it presumably could based on comparison of realized and potential species ranges inferred from niche models (Fig. 5), the latter of which includes the Atlantic coast forest in Brazil and Caribbean islands. It may also explain why, unlike many of the most abundant taxa in lowland rainforests in the northern Neotropics (Bemmels et al. 2018), individual *Otoba* species are restricted to a single side of the Andes. At a local scale, dispersal limitation is observed in *Otoba parvifolia*, whose seeds are dispersed at low frequency, typically over short distances (Terborgh et al. 2011). However, high levels of seed-set, both in closed canopy forests and in treefall gaps, may make *Otoba* an effective colonizer of new habitats once they colonize a new region (Myster 2020).

Despite relatively few major biogeographical movements, rapid niche evolution has allowed *Otoba* to expand into many habitat types in the northern Neotropics. These habitats span a large temperature differential and include areas that are both completely aseasonal (e.g., *O. latialata*) and those with fairly well-defined seasons (e.g., Central American taxa) (Fig. 5; Appendix S9). Rapid climatic niche evolution in *Otoba* facilitates its establishment in new environments upon expansion or dispersal into new habitats, which tend to be in close geographic proximity to ancestral habitats. In fact, the few biogeographic movements that we infer correspond with climatic niche shifts. This includes a shift towards broader temperature range and decreased precipitation seasonality tolerance following dispersal into the western Amazon and towards higher temperature fluctuations in the more seasonal environments of Central America upon expansion via the Isthmus of Panama. The most extreme example of niche evolution in *Otoba* is the sister pair *O. gordoniifolia*, which is restricted to mid-elevations of the Andes and adapted to the coolest temperatures in the genus, and the Chocoan *O. latialata,* which is adapted to the warmest temperatures (Fig. 5). Our comparative analyses demonstrate that movement into a new area is a much stronger constraint on biogeographic evolution in *Otoba* than is the ability to adapt to a new environment upon establishment. There are, however, obvious limits— *Otoba* is not found in high-elevation or very arid habitats, even when they are near their current range. Like many other wet tropical trees (Esquivel-Muelbert et al. 2017), their absence in arid areas is likely due to physiological constraints in the face of drought, while freezing tolerance is an additional important constraint in high elevation grasslands (Koehler, Center, and Cavender-Bares 2012).

## Conclusion

Supporting our central hypothesis within *Otoba*, an ecologically dominant genus from a plant family in which long-distance dispersal is rare to non-existent, adaptation to new environmental niches is much more common than dispersal into pre-adapted environments. Put simply, within physiological constraints related to drought and freezing, it is easier for *Otoba* to evolve than to move. This adds to a growing body of literature demonstrating that dispersal-driven niche evolution following movement from geographically proximal regions is key to the assembly of the Neotropical biota: large portions of its characteristic biomes, including the Amazon (Antonelli et al. 2018) and Cerrado (Simon et al. 2009), are composed of local migrants with ancestors from nearby, but climatically different environments. Notable exceptions are montane environments, including low- to mid-elevation environments where some species of *Otoba* occur (Linan et al. 2021), and high elevation grasslands, where Myristicaceae are absent (Hughes and Eastwood 2006). This points towards an interaction of idiosyncratic, clade-specific traits with steep environmental gradients in driving patterns of plant distribution in the Neotropics.

## Supporting information

Frost et al. Supplemental Material

## Acknowledgements

This research was funded by a Louisiana Board of Regents Research Competitiveness Subprogram grant and by the LSU Department of Biological Sciences. We would like to thank the Missouri Botanical Garden (MO) for their access to their important collections, and Reinaldo Aguilar, Rudy Gelis, Timothy Paine, and John Janovec for permission to use their photographs. We thank Brant Faircloth, Matthew Johnson, Carl Oliveros, and Jessie Salter for their guidance in library preparation and Brant Faircloth for access to laboratory equipment, and LSU HPC for computational resources. This manuscript benefited from feedback from Laymon Ball, Janet Mansaray, and Diego Paredes-Burneo.

## Data Availability Statement

Illumina reads will be submitted to the NCBI Sequence Read Archive (SRA) and all other data formats (tree files, alignments, character matrices, etc.) will be uploaded on the Dryad Digital Repository and made available upon publication.

## Biosketch

The Lagomarsino Lab studies Neotropical plant evolution (more at http://www.lauralago.net/). L.F., D.A.S.A, and L.P.L. conceived the ideas; all authors collected data; L.F., D.S., and L.P.L analysed data; and L.F. and L.P.L led the writing.

