## Supplementary material for "Niche evolution of the Neotropical tree genus *Otoba* in the context of global biogeography of the nutmeg family, Myristicaceae": Frost et al. Supplemental Material

*Running title:* Biogeography of *Otoba* and Myristicaceae

Laura Frost, Daniel A. Santamaría-Aguilar, Daisy Singletary, & Laura P. Lagomarsino[1]

Shirley C. Tucker Herbarium, Department of Biological Sciences, Louisiana State University, Baton Rouge, LA 70808, USA

[1]author for communication:

**Appendix S1.** Voucher information for *Otoba* samples, in the following format:

Sample ID; Taxon (Accepted Name); Voucher (Herbarium); Provenance; [Latitude, Longitude].

Otoba_acuminata_MM2720; ***Otoba acuminata*** (Standl.) *A.H. Gentry; Mary Merello, Allison Miller & Beatriz Wong 2720* (MO); Panama; [09°13’10”N, 079°21’38”W]. Otoba_cyclobasis_MT281; ***Otoba cyclobasis*** T.S.Jaram. & Balslev; *Milton Tirado, E. Albuja & M. Chapiro 281* (MO); Ecuador; [00°43’N, 078°53’W]. Otoba_glycycarpa_LV25198; ***Otoba glycycarpa*** (Ducke) W.A.Rodrigues & T.S.Jaram.*; Luis Valenzuela G., Jaime Flores, Gerry Shareva M.,* et al. *25198* (MO); Peru; [11°21’39"S 074°02’31”W]. Otoba_gordoniifolia_RZ196; ***Otoba gordoniifolia*** (A. DC.) A.H. Gentry; *R.A. Zahawi 196* (MO); Ecuador; [00°05’N, 079°40’W]. Otoba_gracilipes_DC884; ***Otoba gracilipes*** (A.C. Sm.) A.H. Gentry; *D. Cárdenas L. 884* (MO); Colombia; [7°26’20.0”N, 77°07’15.8”W]. Otoba_latialata_RC4751; ***Otoba latialata*** (Pittier) A.H. Gentry; *R. Callejas* et al. *4751* (MO); Colombia; [7°0'59.36”N, 76°18’31.52”W]. Otoba_novogranatensis_AG476; ***Otoba novogranatensis*** Moldenke; *A. Grijalva, C. Aulestia & J. Taicúz 476* (MO); Ecuador; [01°02’N, 078°15’W]. Otoba_novogranatensis_CK681; ***Otoba novogranatensis*** Moldenke; *C. Kernan & P. Phillips 681* (MO); Costa Rica; [08°27’N, 083°33’W]. Otoba_novogranatensis_EB500; ***Otoba novogranatensis*** Moldenke; *E. Bello C. 500* (MO); Costa Rica; [10°18’36”N, 084°42’00”W]. Otoba_novogranatensis_GP2325; ***Otoba novogranatensis*** Moldenke; *G. A. Tipaz, P. Méndez, H. Vargas & M. Chapiro 2325* (MO); Ecuador; [00°45’N, 078°47’W]. Otoba_novogranatensis_LG20482; ***Otoba novogranatensis*** Moldenke; *L. D. Gómez P., R. L. Liesner & E. J. Judziewicz 20482* (MO); Costa Rica; [09°38’24”N, 082°48’30”W]. Otoba_novogranatensis_WP16081; ***Otoba novogranatensis*** Moldenke; *W. A. Palacios & M. Tirado 16081* (MO); Ecuador; [00°45’N, 078°56’W]. Otoba_novogranatensis_WS36336; ***Otoba novogranatensis*** Moldenke; *W. D. Stevens & O. M. Montiel J. 36336* (MO); Nicaragua; [12°17’36”N, 085°05’58”W]. Otoba_parvifolia_DN9151; ***Otoba parvifolia*** (Markgr.) *A.H.Gentry; D. A. Neill, F. Hurtado & A. A. Alvarado 9151* (MO); Ecuador, Napo; [00°36’S, 077°23’W]. Otoba_parvifolia_MN37243; ***Otoba parvifolia*** (Markgr.) A.H.Gentry; *M. H. Nee 37243* (MO); Bolivia; [17°39’S, 063°43’W] Otoba_parvifolia_MS1182; ***Otoba parvifolia*** (Markgr.) A.H.Gentry; *M.S. Silveira 1182* (MO); Brazil, Acre; [08°17’48”S, 071°08’36”W]. Otoba_parvifolia_RV19070; ***Otoba parvifolia*** (Markgr.) A.H.Gentry; *R. Vásquez & R. Apanú 19070* (MO); 08 September 1994; Peru, Amazonas, Condorcanqui; 320 m; [04°51’S, 078°18’W]. Otoba_sp_nov_JP16902; **n/a**; *J. J. Pipoly, III, Á. Cogollo P.* et al. *16902* (MO); Colombia; [06°29’N, 076°14’W]. Otoba_sp_nov_RC5752; **n/a**; *R. Callejas, R. Fonnegra G., F. J. Roldán & A. L. Arbeláez 5752* (MO); Colombia; [07°20’N, 076°30’W]. Otoba_vespertilio_GM12543; ***Otoba vespertilio*** D. Santam. & J.E. Jiménez; *Gordon McPherson 12543* (MO); Panama; [08°47’03”N, 082°10’52”W].

**Appendix S2.** Myristicaceae GenBank accession numbers

| Species | matK | rbcL | ndhF |
|---|---|---|---|
| *Ambavia gerrardii* | AY220435 | - | AY218168 |
| *Anaxagorea acuminata* | AY220436 | - | AY218169 |
| *Annickia kummeriae* | AY238961 | - | - |
| *Annona muricata* | - | - | AY218170 |
| *Artabotrys hexapetalus* | AY238962 | - | - |
| *Asimina triloba* | AY220437 | - | AY218171 |
| *Brochoneura acuminata* | AY220442 | - | AY218179 |
| *Cananga odorata* | AY220438 | - | AY218172 |
| *Cassytha filiformis* | - | - | - |
| *Cephalosphaera usambarensis* | AY220443 | - | AY218180 |
| *Coelocaryon oxycarpum* | AY220444 | - | AY218181 |
| *Compsoneura atopa* | EU090469 | EU090508 | - |
| *Compsoneura capitellata* | EU090473 | EU090509 | - |
| *Compsoneura debilis* | EU090475 | EU090514 | - |
| *Compsoneura excelsa* | EU090482 | EU090518 | - |
| *Compsoneura mutisii* | EU090496 | EU090532 | - |
| *Compsoneura sprucei* | AY220445 | EU090539 | AY218182 |
| *Compsoneura ulei* | EU090505 | EU090541 | - |
| *Degeneria roseiflora* | AY220440 | - | AY218174 |
| *Gymnacranthera contracta* | MH332593 | - | - |
| *Gymnacranthera forbesii* | MH332622 | - | - |
| *Horsfieldia amygdalina* | KR530941 | KR529438 | - |
| *Horsfieldia amygdalina* | MF547527 | - | - |
| *Horsfieldia grandis* | MH332620 | - | - |
| *Horsfieldia kingii* | KR530945 | KR529441 | - |
| *Horsfieldia pandurifolia* | NC042225 | NC042225 | NC042225 |
| *Horsfieldia polyspherula* | KJ708961 | MG817045 | - |
| *Horsfieldia prainii* | KR530951 | KR529444 | - |
| *Horsfieldia pulcherrima* | MG816907 | MG817055 | - |
| *Horsfieldia punctatifolia* | AY220448 | - | AY218184 |
| *Horsfieldia subglobosa* | MH332591 | - | - |
| *Iryanthera hostmanni* | AY220449 | JQ625774 | AY218185 |
| *Iryanthera sagotiana* | FJ514645 | JQ625975 | - |
| *Isolona campanulata* | AY238963 | - | - |
| *Knema cinerea* | KJ708967 | KJ594758 | - |
| *Knema elegans* | KR530962 | KR529456 | - |
| *Knema furfuracea* | KR530963 | KR529457 | - |
| *Knema globularia* | AB924868 | KR529464 | - |
| *Knema hookeriana* | KJ708968 | KJ594760 | - |
| *Knema laurina* | AY220450 | KJ594761 | AY218186 |
| *Knema lenta* | KR530973 | KR529465 | - |

| | | | |
|---|---|---|---|
| ***Knema linifolia*** | **KR530976** | **KR529470** | **-** |
| ***Knema patentinervia*** | **KJ708971** | **KJ594762** | **-** |
| ***Knema tenuinervia*** | **KR530977** | **KR529476** | **-** |
| ***Malmea dielsiana*** | **AY238964** | **AY238955** | **-** |
| ***Mauloutchia chapelieri*** | **AY220451** | **AF197594** | **AY218187** |
| ***Michelia maudiae*** | **MN990580** | **MN990580** | **MN990580** |
| ***Myristica andamanica*** | **MF547528** | **MF158638** | **-** |
| ***Myristica beddomei*** | **MF547537** | **MF186600** | **-** |
| ***Myristica cinnamomea*** | **KJ709010** | **KJ594811** | **-** |
| ***Myristica fragrans*** | **KT445278** | **AF206798** | **AY218188** |
| ***Myristica gigantea*** | **MG816899** | **MG817047** | **-** |
| ***Myristica malabarica*** | **MF547530** | **KY945260** | **-** |
| ***Osteophloeum platyspermum*** | **JQ626371** | **JQ625884** | **-** |
| ***Polyalthia suberosa*** | **AY220439** | **-** | **-** |
| ***Pycnanthus angolensis*** | **AY220453** | **-** | **AY218189** |
| ***Staudtia gabonensis*** | **KC627785** | **KC628454** | **-** |
| ***Staudtia kamerunensis*** | **KC627748** | **KC628429** | **-** |
| ***Uvaria afzelii*** | **AY238966** | **-** | **-** |
| ***Virola koschnyi*** | **EU669473** | **JQ592895** | **-** |
| ***Virola kwatae*** | **FJ514688** | **JQ626043** | **-** |
| ***Virola multiflora*** | **GQ982125** | **GQ981913** | **-** |
| ***Virola nobilis*** | **GQ982126** | **GQ981914** | **-** |
| ***Xylopia peruviana*** | **AY238967** | **-** | **-** |

**Appendix S3.** Summary of sequence data recovered for each sample for targeted and off-target (cpDNA) loci. Asterisks at sample names indicate samples that were not included in any phylogenetic analysis. Samples for which there was insufficient cpDNA data to include in phylogenetic analyses are marked with “n/a”.

| Sample | Mean sequence length (bp) | # of loci | Length of ungapped cpDNA (bp) | Collection year | Annual precipitation (mm) |
|---|---|---|---|---|---|
| ***Otoba acuminata*_MM2720** | 209.25 | 4 | 39,553 | 2001 | 3,021 |
| ***Otoba cyclobasis*_MT281** | 183.9 | 73 | 3,399 | 1993 | 2,265 |
| ***Otoba glycycarpa*_LV25198** | 227.88 | 96 | 32,401 | 2013 | 1,842 |
| ***Otoba gordoniifolia*_RZ196** | 206.51 | 61 | 20,318 | 1996 | 1,933 |
| **Otoba_gracilipes_DC884*** | 192 | 2 | n/a | 1987 | 3,065 |
| ***Otoba latialata*_RC4751** | 244.39 | 142 | 42,374 | 1987 | 2,610 |
| ***Otoba novogranatensis*_AG476** | 102 | 1 | 3,533 | 1993 | 3,048 |
| ***Otoba novogranatensis*_CK681** | 174 | 2 | 13,653 | 1988 | 3,499 |
| ***Otoba novogranatensis*_EB500*** | 208.91 | 11 | n/a | 1988 | 3,524 |
| ***Otoba novogranatensis*_GP2325** | 186.75 | 4 | 41,378 | 1992 | 2,057 |
| ***Otoba novogranatensis*_LG20482** | 299.92 | 157 | 111,863 | 1983 | 3,244 |
| ***Otoba novogranatensis*_WP16081** | 90 | 1 | 47,902 | 1993 | 2,326 |
| ***Otoba novogranatensis*_WS36336** | 457.56 | 217 | n/a | 2015 | 1,864 |
| ***Otoba parvifolia*_DN9151*** | 183 | 4 | n/a | 1989 | 3,870 |
| ***Otoba parvifolia*_MN37243** | 173.47 | 17 | 28,315 | 1988 | 1,449 |
| ***Otoba parvifolia*_MS1182** | 139.5 | 6 | 72,683 | 1995 | 2,109 |
| ***Otoba parvifolia*_RV19070** | 215.62 | 86 | n/a | 1994 | 2,095 |
| ***Otoba sp. nov.*_JP16902*** | 145.41 | 17 | n/a | 1992 | 2,438 |
| ***Otoba sp. nov*_RC5752*** | 126 | 5 | n/a | 1987 | 3,732 |
| ***Otoba vespertilio*_GM12543** | 131 | 3 | 1,532 | 1988 | 3,561 |

**Appendix S4**. Heatmap showing the percent reference protein length recovered for all samples across all 353 loci.

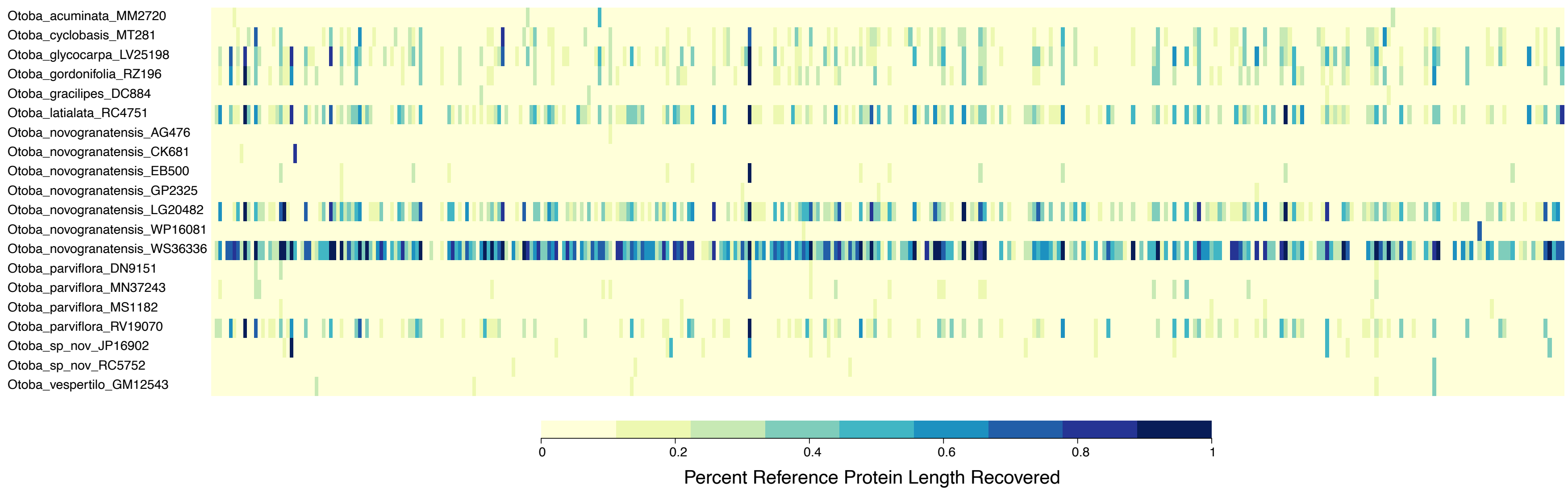

**Appendix S5.** Divergence date estimations of *Otoba* using difference calibration schema (denoted to left of phylogeny).

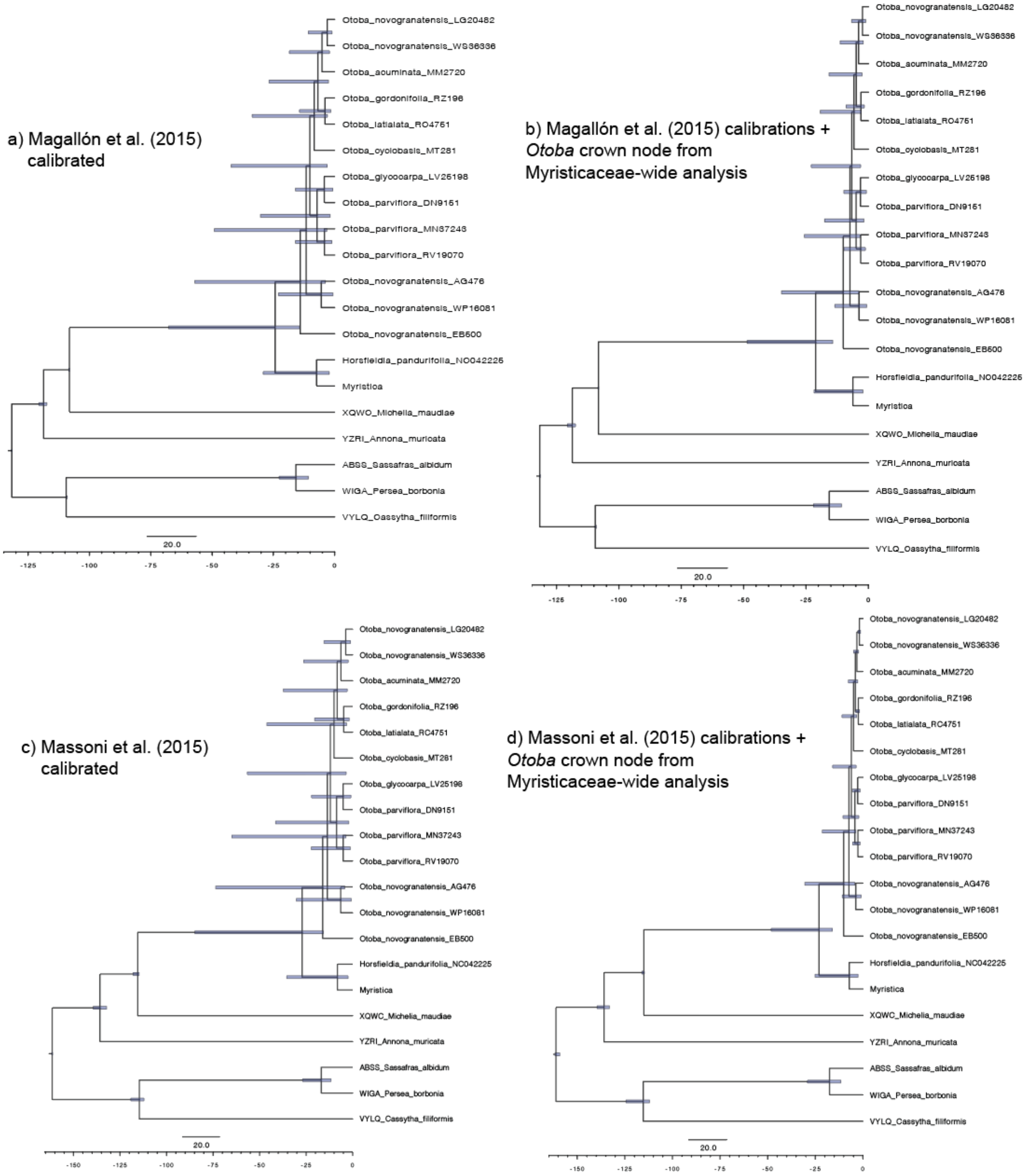


**Figure S5.** treePL divergence date estimations with different calibration schemes. a) and b) include secondary calibrations derived from Magallón et al. (2015) at the nodes corresponding to Laurales + Magnoliales, Laurales, Magnoliales, and Myristicaceae; b) additionally includes a calibration for the crown node of Otoba derived from our densely sampled, family-wide BEAST2 analysis of Myristicaceae calibrated with dates from Magallón et al (2015). c) and d) include secondary calibrations derived from Massoni et al. (2015) at the nodes corresponding to Laurales + Magnoliales, Laurales, Magnoliales, and Myristiaceae; d) additionally includes a calibration for the crown node of Otoba derived from our densely sampled, family-wide BEAST2 analysis of Myristicaceae calibrated with dates from Massoni et al (2015). Bars at nodes represent 95% confidence intervals as inferred across 100 RAxML bootstrap replicates.

**Appendix S6.** Divergence date estimation from BEAST2 analysis of Myristiaceae using the Magallón et al. (2015) calibration scheme. Values at nodes correspond to the posterior probability, while bars at nodes represented the 95% HPD on the estimated age of the node, with time scale below phylogeny in millions of years before present. (Figure on next page)

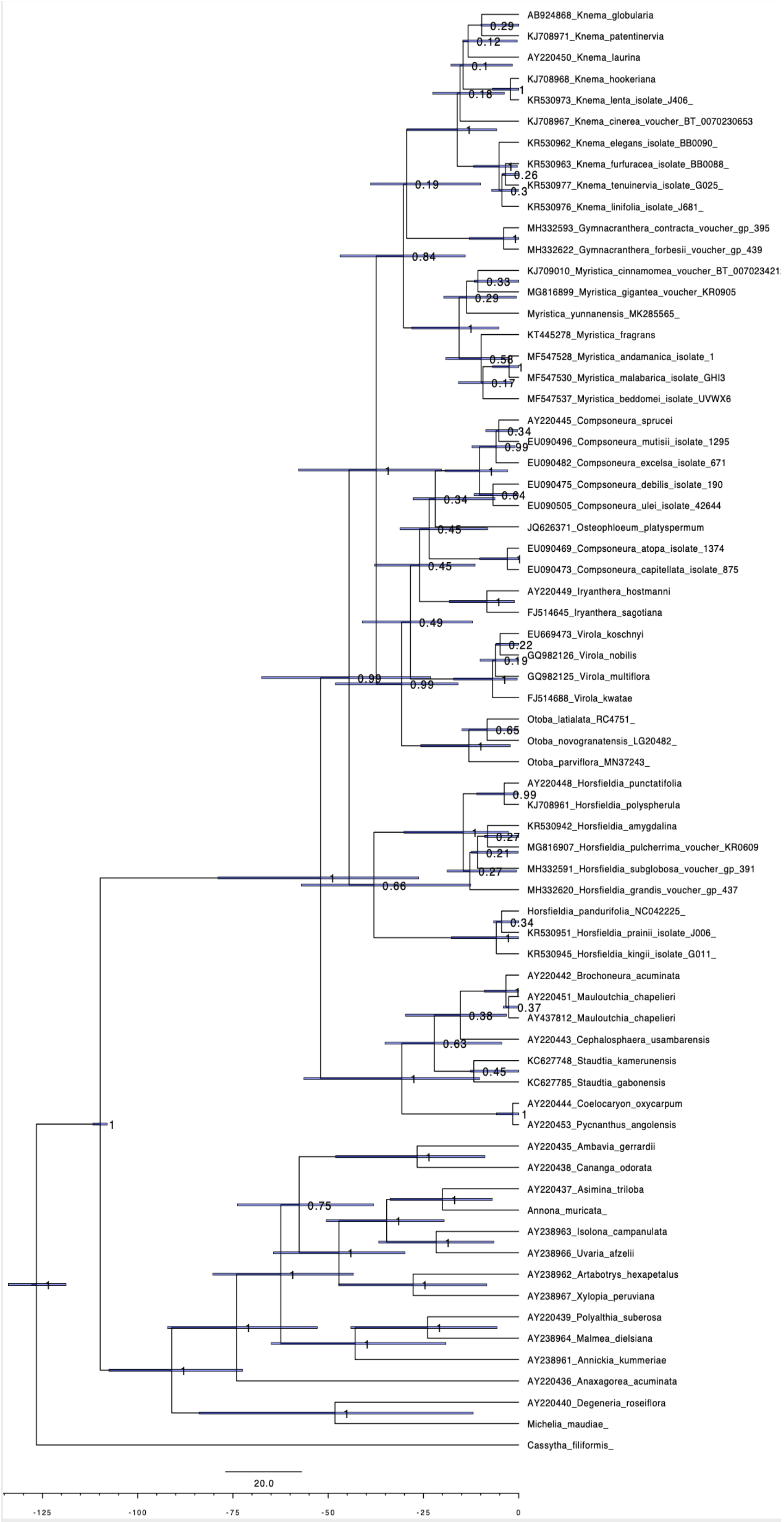
AB924868_Knema_globularia
KJ708971_Knema_patentinervia
AY220450_Knema_laurina
KJ708968_Knema_hookeriana
KR530973_Knema_lenta_isolate_J406_
KJ708967_Knema_cinerea_voucher_BT_0070230653
KR530962_Knema_elegans_isolate_BB0090_
KR530963_Knema_furfuracea_isolate_BB0088_
KR530977_Knema_tenuinervia_isolate_G025_
KR530976_Knema_linifolia_isolate_J681_
MH332593_Gymnacranthera_contracta_voucher_gp_395
MH332622_Gymnacranthera_forbesii_voucher_gp_439
MG816899_Myristica_gigantea_voucher_KR0905
Myristica_yunnanensis_MK285565_
KT445278_Myristica_fragrans
MF547528_Myristica_andamanica_isolate_1
MF547530_Myristica_malabarica_isolate_GHI3
MF547537_Myristica_beddomei_isolate_UVWX6
AY220445_Compsoneura_sprucei
EU090496_Compsoneura_mutisii_isolate_1295
EU090482_Compsoneura_excelsa_isolate_671
EU090475_Compsoneura_debilis_isolate_190
EU090505_Compsoneura_ulei_isolate_42644
JQ626371_Osteophloeum_platyspermum
EU090469_Compsoneura_atopa_isolate_1374
EU090473_Compsoneura_capitellata_isolate_875
AY220449_Iryanthera_hostmanni
FJ514645_Iryanthera_sagotiana
EU669473_Virola_koschnyi
GQ982126_Virola_nobilis
GQ982125_Virola_multiflora
FJ514688_Virola_kwatae
Otoba_latialata_RC4751_
Otoba_novogranatensis_LG20482_
Otoba_parviflora_MN37243_
AY220448_Horsfieldia_punctatifolia
KJ708961_Horsfieldia_polyspherula
KR530942_Horsfieldia_amygdalina
MG816907_Horsfieldia_pulcherrima_voucher_KR0609
MH332591_Horsfieldia_subglobosa_voucher_gp_391
MH332620_Horsfieldia_grandis_voucher_gp_437
Horsfieldia_pandurifolia_NC042225_
KR530951_Horsfieldia_prainii_isolate_J006_
KR530945_Horsfieldia_kingii_isolate_G011_
AY220442_Brochoneura_acuminata
AY220451_Mauloutchia_chapelieri
AY437812_Mauloutchia_chapelieri
AY220443_Cephalosphaera_usambarensis
KC627748_Staudtia_kamerunensis
KC627785_Staudtia_gabonensis
AY220444_Coelocaryon_oxycarpum
AY220453_Pycnanthus_angolensis
AY220435_Ambavia_gerrardii
AY220438_Cananga_odorata
AY220437_Asimina_triloba
Annona_muricata_
AY238963_Isolona_campanulata
AY238966_Uvaria_afzelii
AY238962_Artabotrys_hexapetalus
AY238967_Xylopia_peruviana
AY220439_Polyalthia_suberosa
AY238964_Malmea_dielsiana
AY238961_Annickia_kummeriae
AY220436_Anaxagorea_acuminata
AY220440_Degeneria_roseiflora
Michelia_maudiae_
Cassytha_filiformis_
20.0
-125
-100
-75
-50
-25
0

**Appendix S7.** Divergence date estimation from BEAST2 analysis of Myristicaceae using the Massoni et al. (2015) calibration scheme. Values at nodes correspond to the posterior probability, while bars at nodes represented the 95% HPD on the estimated age of the node with time scale below phylogeny in millions of years before present. (Figure on next page)

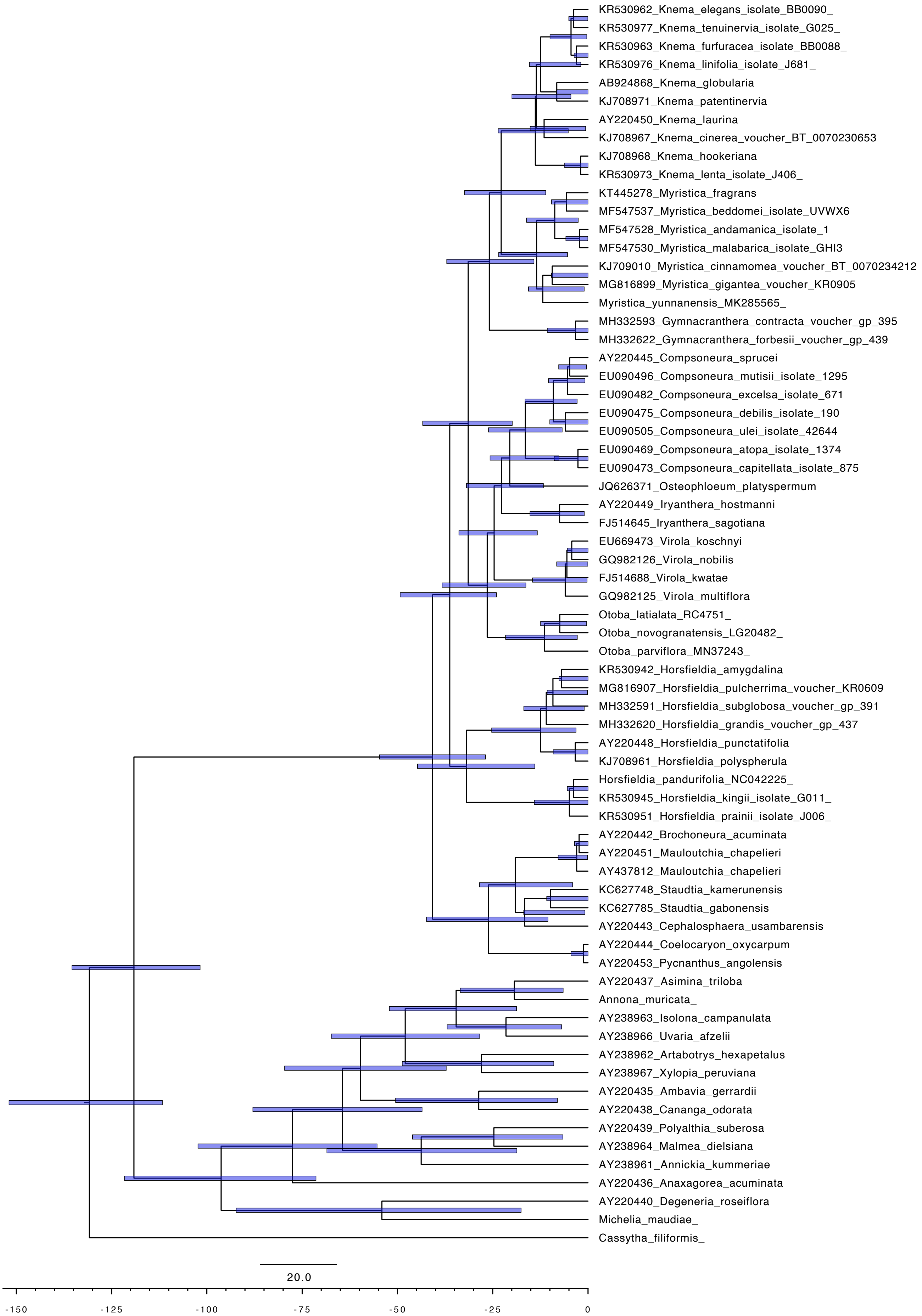
KR530962_Knema_elegans_isolate_BB0090_
KR530977_Knema_tenuinervia_isolate_G025_
KR530963_Knema_furfuracea_isolate_BB0088_
KR530976_Knema_linifolia_isolate_J681_
AB924868_Knema_globularia
KJ708971_Knema_patentinervia
AY220450_Knema_laurina
KJ708967_Knema_cinerea_voucher_BT_0070230653
KJ708968_Knema_hookeriana
KR530973_Knema_lenta_isolate_J406_
KT445278_Myristica_fragrans
MF547537_Myristica_beddomei_isolate_UVWX6
MF547528_Myristica_andamanica_isolate_1
MF547530_Myristica_malabarica_isolate_GHI3
KJ709010_Myristica_cinnamomea_voucher_BT_0070234212
MG816899_Myristica_gigantea_voucher_KR0905
Myristica_yunnanensis_MK285565_
MH332593_Gymnacranthera_contracta_voucher_gp_395
MH332622_Gymnacranthera_forbesii_voucher_gp_439
AY220445_Compsoneura_sprucei
EU090496_Compsoneura_mutisii_isolate_1295
EU090482_Compsoneura_excelsa_isolate_671
EU090475_Compsoneura_debilis_isolate_190
EU090505_Compsoneura_ulei_isolate_42644
EU090469_Compsoneura_atopa_isolate_1374
EU090473_Compsoneura_capitellata_isolate_875
JQ626371_Osteophloeum_platyspermum
AY220449_Iryanthera_hostmanni
FJ514645_Iryanthera_sagotiana
EU669473_Virola_koschnyi
GQ982126_Virola_nobilis
FJ514688_Virola_kwatae
GQ982125_Virola_multiflora
Otoba_latialata_RC4751_
Otoba_novogranatensis_LG20482_
Otoba_parviflora_MN37243_
KR530942_Horsfieldia_amygdalina
MG816907_Horsfieldia_pulcherrima_voucher_KR0609
MH332591_Horsfieldia_subglobosa_voucher_gp_391
MH332620_Horsfieldia_grandis_voucher_gp_437
AY220448_Horsfieldia_punctatifolia
KJ708961_Horsfieldia_polyspherula
Horsfieldia_pandurifolia_NC042225_
KR530945_Horsfieldia_kingii_isolate_G011_
KR530951_Horsfieldia_prainii_isolate_J006_
AY220442_Brochoneura_acuminata
AY220451_Mauloutchia_chapelieri
AY437812_Mauloutchia_chapelieri
KC627748_Staudtia_kamerunensis
KC627785_Staudtia_gabonensis
AY220443_Cephalosphaera_usambarensis
AY220444_Coelocaryon_oxycarpum
AY220453_Pycnanthus_angolensis
AY220437_Asimina_triloba
Annona_muricata_
AY238963_Isolona_campanulata
AY238966_Uvaria_afzelii
AY238962_Artabotrys_hexapetalus
AY238967_Xylopia_peruviana
AY220435_Ambavia_gerrardii
AY220438_Cananga_odorata
AY220439_Polyalthia_suberosa
AY238964_Malmea_dielsiana
AY238961_Annickia_kummeriae
AY220436_Anaxagorea_acuminata
AY220440_Degeneria_roseiflora
Michelia_maudiae_
Cassytha_filiformis_
20.0
-150
-125
-100
-75
-50
-25
0

**Appendix S8.** Average values for each BIOCLIM variable and elevation in *Otoba*. A description of each bioclimatic variable can be found on the WorldClim website (https://www.worldclim.org/data/bioclim.html).

| species | BIO1 | BIO2 | BIO3 | BIO4 | BIO5 | BIO6 | BIO7 | BIO8 | BIO9 | BIO10 | BIO11 | BIO12 | BIO13 | BIO14 | BIO15 | BIO16 | BIO17 | BIO18 | BIO19 | Elevation |
|---|---|---|---|---|---|---|---|---|---|---|---|---|---|---|---|---|---|---|---|---|
| acuminata | 24.64 | 8.05 | 81.37 | 59.04 | 29.81 | 19.92 | 9.89 | 24.43 | 24.92 | 25.45 | 23.99 | 2917.25 | 413.34 | 66.94 | 48.61 | 1066.25 | 275.81 | 522.28 | 852.25 | 404.84 |
| cyclobasis | 23.72 | 8.52 | 87.41 | 33.67 | 28.73 | 18.97 | 9.76 | 24.08 | 23.48 | 24.17 | 23.38 | 2224.57 | 283.14 | 100.43 | 35.38 | 794.71 | 356.86 | 741.43 | 429.71 | 405.86 |
| glycycarpa | 24.04 | 9.53 | 82.77 | 50.71 | 29.60 | 18.08 | 11.51 | 23.93 | 23.97 | 24.54 | 23.32 | 2927.28 | 337.47 | 162.28 | 26.65 | 952.24 | 515.58 | 682.83 | 714.08 | 543.70 |
| gordoniifolia | 19.46 | 8.93 | 88.10 | 36.04 | 24.58 | 14.42 | 10.15 | 19.52 | 19.40 | 19.89 | 19.04 | 2067.50 | 296.54 | 64.08 | 49.44 | 814.19 | 245.00 | 676.53 | 473.68 | 1380.33 |
| latialata | 24.55 | 8.17 | 86.77 | 38.33 | 29.37 | 19.96 | 9.40 | 24.23 | 24.73 | 25.04 | 24.09 | 4127.77 | 522.05 | 168.54 | 37.46 | 1392.22 | 628.37 | 868.07 | 1248.70 | 448.46 |
| novogranatensisCA | 24.20 | 8.67 | 80.03 | 69.02 | 29.78 | 18.96 | 10.82 | 23.92 | 24.42 | 25.15 | 23.42 | 3031.06 | 473.82 | 46.26 | 56.74 | 1213.22 | 209.37 | 537.39 | 885.85 | 432.73 |
| novogranatensisSA | 22.52 | 8.67 | 87.29 | 39.20 | 27.55 | 17.60 | 9.95 | 22.62 | 22.43 | 23.00 | 22.05 | 2752.89 | 367.19 | 101.41 | 44.34 | 1003.30 | 371.61 | 774.67 | 692.98 | 799.55 |
| parvifolia | 24.18 | 9.79 | 82.33 | 60.19 | 29.90 | 17.97 | 11.93 | 24.25 | 23.85 | 24.77 | 23.33 | 2710.19 | 325.99 | 135.41 | 32.68 | 918.32 | 438.74 | 677.39 | 593.91 | 511.48 |

**Appendix S9.** Loadings of phylogenetic PCA of BIOCLIM variables in *Otoba*. A description of each bioclimatic variable can be found on the WorldClim website (*https://www.worldclim.org/data/bioclim.html*).

| | PC1 | PC2 | PC3 | PC4 |
|---|---|---|---|---|
| **BIO1** | 0.9590468 | 0.14849091 | -0.12600397 | 0.20116525 |
| **BIO2** | -0.4602410 | 0.20067826 | -0.77533178 | -0.36089120 |
| **BIO3** | -0.4027127 | -0.89055722 | 0.13301025 | 0.08391671 |
| **BIO4** | 0.2890433 | 0.92875105 | -0.08879483 | -0.19983694 |
| **BIO5** | 0.9271045 | 0.26345593 | -0.20576943 | 0.15920598 |
| **BIO6** | 0.9564835 | 0.03836619 | 0.07135413 | 0.27848748 |
| **BIO7** | -0.2007299 | 0.56990207 | -0.71722923 | -0.34127397 |
| **BIO8** | 0.9319273 | 0.12154685 | -0.17278971 | 0.28132250 |
| **BIO9** | 0.9685474 | 0.17064102 | -0.05535913 | 0.17190043 |
| **BIO10** | 0.9502414 | 0.22187517 | -0.09810446 | 0.18980573 |
| **BIO11** | 0.9617379 | 0.08309425 | -0.09836778 | 0.23662162 |
| **BIO12** | 0.9104890 | -0.15241647 | 0.15008476 | -0.35080041 |
| **BIO13** | 0.8240385 | 0.09699091 | 0.38962441 | -0.39547895 |
| **BIO14** | 0.6495930 | -0.58925208 | -0.42578738 | -0.18673285 |
| **BIO15** | -0.3713011 | 0.56788707 | 0.69624654 | -0.16370803 |
| **BIO16** | 0.8466405 | 0.01969772 | 0.32392806 | -0.41709562 |
| **BIO17** | 0.7224432 | -0.59367026 | -0.27452234 | -0.20035213 |
| **BIO18** | 0.3329619 | -0.88486442 | -0.09434808 | -0.17350519 |
| **BIO19** | 0.8706295 | -0.06163663 | 0.29534788 | -0.37339337 |

**Appendix S10.** Ancestral state reconstruction of individual phylogenetic principal components of BIOCLIM variables in *Otoba* (See Appendix S9). Analysis conducted with contMap in phytools.

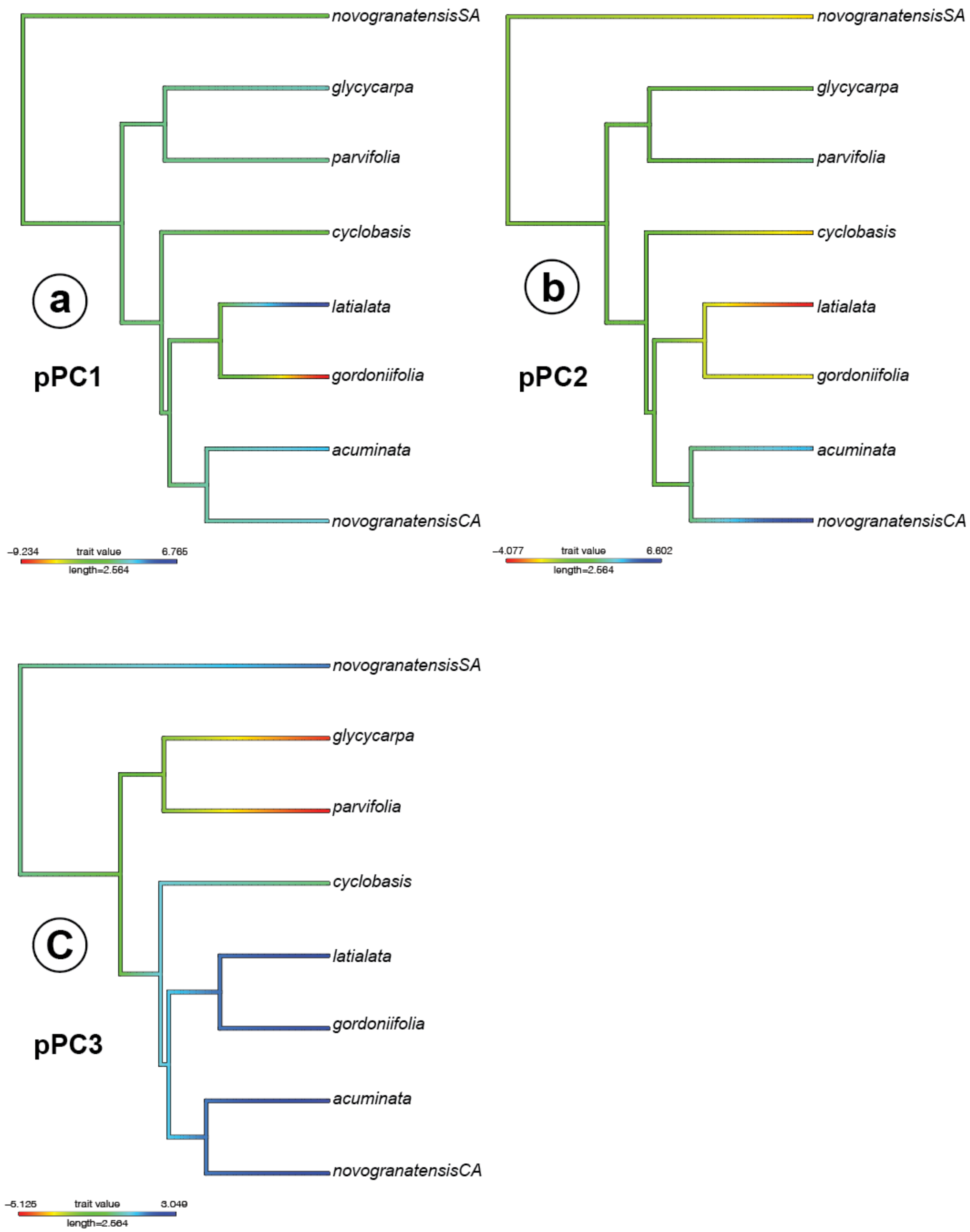
